# Satellite Imaging of Global Urbanicity relate to Adolescent Brain Development and Behavior

**DOI:** 10.1101/781674

**Authors:** Jiayuan Xu, Xiaoxuan Liu, Alex Ing, Qiaojun Li, Wen Qin, Lining Guo, Conghong Huang, Jingliang Cheng, Meiyun Wang, Zuojun Geng, Wenzhen Zhu, Bing Zhang, Weihua Liao, Shijun Qiu, Hui Zhang, Xiaojun Xu, Yongqiang Yu, Bo Gao, Tong Han, Guangbin Cui, Feng Chen, Junfang Xian, Jiance Li, Jing Zhang, Xinian Zuo, Dawei Wang, Wen Shen, Yanwei Miao, Fei Yuan, Su Lui, Xiaochu Zhang, Kai Xu, Longjiang Zhang, Zhaoxiang Ye, Tobias Banaschewski, Gareth J. Barker, Arun L.W. Bokde, Erin Burke Quinlan, Sylvane Desrivières, Herta Flor, Antoine Grigis, Hugh Garavan, Penny Gowland, Andreas Heinz, Rüdiger Brühl, Jean-Luc Martinot, Eric Artiges, Frauke Nees, Dimitri Papadopoulos Orfanos, Herve Lemaitre, Tomáš Paus, Luise Poustka, Sarah Hohmann, Juliane H. Fröhner, Michael N. Smolka, Henrik Walter, Robert Whelan, Ran Goldblatt, Kevin Patrick, Vince Calhoun, Mulin JunLi, Meng Liang, Peng Gong, Edward D Barker, Nicholas Clinton, Le Yu, Chunshui Yu, Gunter Schumann, the CHIMGEN and IMAGEN Consortia

**Affiliations:** Department of Radiology and Tianjin Key Laboratory of Functional Imaging, Tianjin Medical University General Hospital, Tianjin 300052, P.R. China; Centre for Population Neuroscience and Precision Medicine (PONS), Institute of Psychiatry, Psychology and Neuroscience, SGDP Centre, King’s College London, London SE5 8AF, United Kingdom; Ministry of Education Key Laboratory for Earth System Modeling, Department of Earth System Science, Tsinghua University, Beijing 100084, P.R. China; Joint Center for Global Change Studies, Beijing 100875, P.R. China; College of Information Engineering, Tianjin University of Commerce, Tianjin 300052, P.R. China; Department of Magnetic Resonance Imaging, The First Affiliated Hospital of Zhengzhou University,450052 Zhengzhou, China; Department of Radiology, Zhengzhou University People’s Hospital and Henan Provincial People’s Hospital, 450003 Zhengzhou, China; Department of Medical Imaging, The Second Hospital of Hebei Medical University, 050000 Shijiazhuang, China; Department of Radiology, Tongji Hospital, Tongji Medical College, Huazhong University of Science and Technology, 430030 Wuhan, China; Department of Radiology, Drum Tower Hospital, Medical School of Nanjing University, 210008 Nanjing, China; Department of Radiology, Xiangya Hospital, Central South University, 410008 Changsha, China; Department of Medical Imaging, The First Affiliated Hospital of Guangzhou University of Chinese Medicine, 510405 Guangzhou, China; Department of Radiology, The First Hospital of Shanxi Medical University, 030001 Taiyuan, China; Department of Radiology, The Second Affiliated Hospital of Zhejiang University, School of Medicine, 310009 Hangzhou, China; Department of Radiology, The First Affiliated Hospital of Anhui Medical University, 230022 Hefei, China; Department of Radiology, Yantai Yuhuangding Hospital, 264000 Yantai, China; Department of Radiology, Tianjin Huanhu Hospital, 300350 Tianjin, China; Functional and Molecular Imaging Key Lab of Shaanxi Province & Department of Radiology, Tangdu Hospital, the Military Medical University of PLA Airforce (Fourth Military Medical University), 710038 Xi’an, China; Department of Radiology, Hainan General Hospital, 570311 Haikou, China; Department of Radiology, Beijing Tongren Hospital, Capital Medical University, 100730 Beijing, China; Department of Radiology, The First Affiliated Hospital of Wenzhou Medical University, 325000 Wenzhou, China; Department of Magnetic Resonance, Lanzhou University Second Hospital, 730050 Lanzhou, China; Department of Psychology, University of Chinese Academy of Sciences (CAS), 100049 Beijing, China; Department of Radiology, Qilu Hospital of Shandong University, 250012 Jinan, China; Department of Radiology, Tianjin First Center Hospital, 300192 Tianjin, China; Department of Radiology, The First Affiliated Hospital of Dalian Medical University, 116011 Dalian, China; Department of Radiology, Pingjin Hospital, Logistics University of Chinese People’s Armed Police Forces, 300162 Tianjin, China; Department of Radiology, the Center for Medical Imaging, West China Hospital of Sichuan University, 610041 Chengdu, China; School of Life Sciences, University of Science & Technology of China, 230026 Hefei, China; Department of Radiology, The Affiliated Hospital of Xuzhou Medical University, 221006 Xuzhou, China; Department of Medical Imaging, Jinling Hospital, Medical School of Nanjing University,210002 Nanjing, China; Department of Radiology, Tianjin Medical University Cancer Institute and Hospital, 300060 Tianjin, China; Department of Child and Adolescent Psychiatry and Psychotherapy, Central Institute of Mental Health, Medical Faculty Mannheim, Heidelberg University, Square J5, 68159 Mannheim, Germany; Department of Neuroimaging, Institute of Psychiatry, Psychology & Neuroscience, King’s College London, United Kingdom; Discipline of Psychiatry, School of Medicine and Trinity College Institute of Neuroscience, Trinity College Dublin, Dublin, Ireland; Department of Cognitive and Clinical Neuroscience, Central Institute of Mental Health, Medical Faculty Mannheim, Heidelberg University, Square J5, Mannheim, Germany; Department of Psychology, School of Social Sciences, University of Mannheim, 68131 Mannheim, Germany; NeuroSpin, CEA, Université Paris-Saclay, F-91191 Gif-sur-Yvette, France; Departments of Psychiatry and Psychology, University of Vermont, 05405 Burlington, Vermont, USA; Sir Peter Mansfield Imaging Centre School of Physics and Astronomy, University of Nottingham, University Park, Nottingham, United Kingdom; Charité – Universitätsmedizin Berlin, corporate member of Freie Universität Berlin, Humboldt-Universität zu Berlin, and Berlin Institute of Health, Department of Psychiatry and Psychotherapy, Campus Charité Mitte, Charitéplatz 1, Berlin, Germany; Physikalisch-Technische Bundesanstalt (PTB), Abbestr. 2 - 12, Berlin, Germany; Institut National de la Santé et de la Recherche Médicale, INSERM Unit 1000 “Neuroimaging & Psychiatry”, University Paris Sud, University Paris Descartes - Sorbonne Paris Cité; and Maison de Solenn, Paris, France; Institut National de la Santé et de la Recherche Médicale, INSERM Unit 1000 “Neuroimaging & Psychiatry”, University Paris Sud, University Paris Descartes - Sorbonne Paris Cité; and Psychiatry Department 91G16, Orsay Hospital, France; Institut National de la Santé et de la Recherche Médicale, UMR 992 INSERM, CEA, Faculté de médecine, Université Paris-Sud, Université Paris-Saclay, NeuroSpin, F-91191 Gif-sur-Yvette, France; Bloorview Research Institute, Holland Bloorview Kids Rehabilitation Hospital and Departments of Psychology and Psychiatry, University of Toronto, Toronto, Ontario, M6A 2E1, Canada; Department of Child and Adolescent Psychiatry and Psychotherapy, University Medical Centre Göttingen, von-Siebold-Str. 5, 37075, Göttingen, Germany; Department of Psychiatry and Neuroimaging Center, Technische Universität Dresden, Dresden, Germany; School of Psychology and Global Brain Health Institute, Trinity College Dublin, Ireland; School of Global Policy and Strategy, International Lane, University of California San Diego, San Diego, CA 92093; Center for Wireless and Population Health Systems, Department of Family and Preventive Medicine and Calit2’s Qualcomm Institute, University of California, San Diego, 9500 Gilman Drive, Dept. 0811, La Jolla, CA 92093-0811; Tri-institutional Center for Translational Research in Neuroimaging and Data Science (TReNDS) [Georgia State University, Georgia Institute of Technology, Emory University], Atlanta, GA 30303; Collaborative Innovation Center of Tianjin for Medical Epigenetics, Tianjin Key Laboratory of Medical Epigenetics, Department of Pharmacology, Tianjin Medical University, Tianjin 300052, P.R. China.; School of Medical Imaging and Tianjin Key Laboratory of Functional Imaging, Tianjin Medical University, Tianjin 300052, P.R. China; Department of Psychology, Institute of Psychiatry, Psychology and Neuroscience, King’s College London, London SE5 8AF, United Kingdom; Google, Inc., 1600 Amphitheatre Parkway, Mountain View, CA 94043, USA; CAS Center for Excellence in Brain Science and Intelligence Technology, Chinese Academy of Sciences, Shanghai, 200031, P.R. China; Population Neuroscience and Precision Medicine (PONS) LIN-Charite Research Group Dept. of Psychiatry and Psychotherapy, Charite, CCM, Humboldt University, Berlin, Germany and Institute for Science and Technology of Brain-inspired Intelligence (ISTBI), Fudan University, Shanghai, P.R. China.

## Abstract

Urbanicity, the impact of living in urban areas, is among the greatest environmental challenges for mental health. While urbanicity might be distinct in different sociocultural conditions and geographic locations, there are likely to exist common features shared in different areas of the globe. Understanding these common and specific relations of urbanicity with human brain and behavior will enable to assess the impact of urbanicity on mental disorders, especially in childhood and adolescence, where prevention and early interventions are likely to be most effective.

We constructed from satellite-based remote sensing data a factor for urbanicity that was highly correlated with population density ground data. This factor, ‘UrbanSat’ was utilized in the Chinese CHIMGEN sample (N=831) and the longitudinal European IMAGEN cohort (N=810) to investigate if exposure to urbanicity during childhood and adolescence is associated with differences in brain structure and function in young adults, and if these changes are linked to behavior.

Urbanicity was found negatively correlated with medial prefrontal cortex volume and positively correlated with cerebellar vermis volume in young adults from both China and Europe. We found an increased correlation of urbanicity with functional network connectivity within- and between- brain networks in Chinese compared to European participants. Urbanicity was highly correlated with a measure of perceiving a situation from the perspective of others, as well as symptoms of depression in both datasets. These correlations were mediated by the structural and functional brain changes observed. Susceptibility to urbanicity was greatest in two developmental windows during mid-childhood and adolescence.

Using innovative technology, we were able to probe the relationship between urban upbringing with brain change and behavior in different sociocultural conditions and geographic locations. Our findings help to identify shared and distinct determinants of adolescent brain development and mental health in different regions of the world, thus contributing to targeted prevention and early-intervention programs for young people in their unique environment. Our approach may be relevant for public health, policy and urban planning globally.

## Introduction

Mental disorders account for 28% of disease burden among non-communicable diseases^1^. Environmental factors account for up to 50% of the attributable risk for mental disorders^2^. The environmental measures investigated in mental health research are generally limited to individual life events^3^, such as trauma, abuse, neglect, or psychosocial stress. While the impact of these individual life events on brain development and behavior is actively investigated^4^, the influence of the physical environment, and its social consequences, such as urbanization and urbanicity, on mental health is less well studied.

Urbanicity can be defined as the impact of living in urban areas at a given time, and refers to the presence of conditions that are particular to urban areas or present to a much greater extent than in nonurban areas^5^. Urbanicity and urbanization are among the greatest environmental challenges globally. Whereas in 1950, less than 30% of the world’s population lived in urban areas, this number has increased to presently 55% and is expected to rise to 68% in 2050^6^. Moreover, environmental impacts of urbanicity are thought to be the most rapidly growing cause for mental illness^7^. While Europe is among the most stable urbanized regions, Asia is home to 54% of the world’s urban population and subject to massive demographic changes: for example, by 2050 China will have added 255 million urban dwellers^8^. While there are likely to exist common environmental factors related to urbanicity that are shared, there may also be distinct influences in China and Europe that affect people differently and at different times during the lifespan^9^.

As 75% of the lifetime burden from mental disorders emerges by age 20 years^10^, the adverse impact of urbanicity can be best addressed and managed in childhood and adolescence. However, data relating urbanicity to mental health have been mainly ascertained in adults, thus our knowledge on how it affects young people is very limited. For example, urbanization may affect brain function in adults^11^, but there are no such data available for children and adolescents whose continuing brain development renders them particularly susceptible to modifying environmental influences.

To facilitate comparative analyses of urbanicity globally, we require common quantitative and longitudinal environmental measures that are ascertained frequently and are identical across different geographic locations and sociocultural conditions. Remote sensing satellite data, having been continuously recorded since the early 1970’s, provide globally applicable, standardized quantitative environmental measures that enable the continuous tracing of environmental influences going back more than 50 years^12^. Therefore, we were interested in developing satellite data features that are suitable measures for urbanicity in Europe and China. Using these data we then aimed to investigate the relation of urbanicity with structural and functional brain measures in two behavioral neuroimaging datasets of young people: the Chinese CHIMGEN study (www.chimgen.tmu.edu.cn) and the European longitudinal IMAGEN cohort (www.imagen-europe.com)^13^. We tested whether structural and functional brain differences related to urbanicity are similar or distinct in Chinese and European participants, and investigated possible susceptibility windows for the effects of urbanicity during child and adolescent development.

## Results

### Development of a remote sensing satellite measure of urbanicity (UrbanSat)

To develop a satellite-based measure of urbanicity, we carried out a confirmatory factor analysis (CFA) in CHIMGEN (n=3336) and IMAGEN (n=1205), identifying a latent factor derived from measures relevant for urbanicity, including nighttime lights^15^, NDVI^16^, NDBI^17^, NDWI^18^ and global land cover mapping^19^. We then validated this factor, which we termed UrbanSat, by calculating correlations with ground-level population density data from GHSL-POP^14^, which were available for both China and Europe for the years 1990, 2000 and 2015. UrbanSat showed very high correlations with ground-level population grid data in rural, town, city and overall livings, indicating its robustness across time during rapid changes of urbanicity and across countries in Asia and Europe (CHIMGEN: *r*=0.697, *P*=1.76×10^-143^ in 1990; CHIMGEN: *r*=0.712, *P=*4.90×10^-196^ in 2000; *r*=0.722, *P=*5.64×10^-197^ in 2015; IMAGEN: *r*=0.715, *P=*5.27×10^-189^ in 2000-2015; Figure 1).

**Figure 1.**
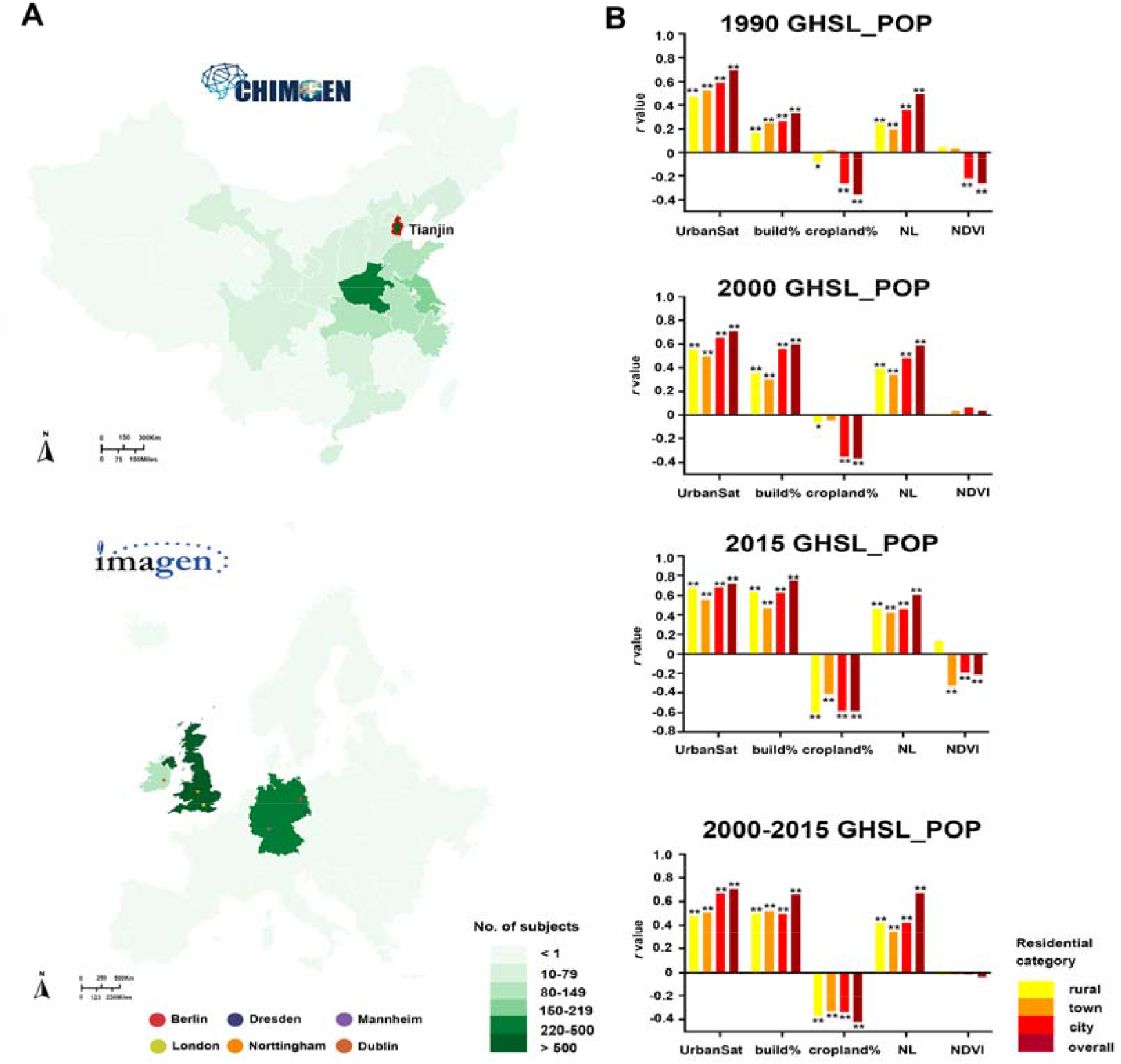
Spatial distribution of subjects in the (A) CHIMGEN (N=3336) and (B) IMAGEN (N=1025) studies. A. The maps showed the numbers of subjects’ distribution from each recruitment center in CHIMGEN and IMAGEN. CHIMGEN Tianjin main center is marked; B. The bars plots show correlations between each satellite feature (UrbanSat, percentage of urban build-up, percentage of cropland, nighttime light and NDVI) and ground-level GHSL population density grid in rural, town, city and overall livings for the years 1990, 2000 and 2015 in CHIMGEN (Top 3). In IMAGEN, we selected only participants who remained at the same address (geoposition) between 2009 and 2015. We therefore used the average population grid between the years 2000 and 2015 (the last one); GHSL_POP, Population density grid of Global Human Settlement Layers; NL, nighttime light. * P < 0.05; ** P < 0.001.

### The relation of UrbanSat with brain gray matter volume and white-matter microstructure

We first tested the correlation of UrbanSat with brain gray matter volume (GMV) in 831 participants from the CHIMGEN (23.81±0.82 years) (Supplementary Table S1) for whom geopositioned data were available since birth. We found a negative correlation of UrbanSat with left medial prefrontal cortex (mPFC) volume (MNI-coordinate: x=-3, y=45, z=27; 4180 voxels; *F* value peak=-6.53; Figure 2A) and a positive correlation with cerebellar vermis volume (MNI-coordinate: x=-4.5, y=-58.5, z=-10.5; 678 voxels; *F* value peak=5.71; Figure 2A) (FWE-correction, voxel *P*<0.05, cluster-size>100 voxels).

**Figure 2.**
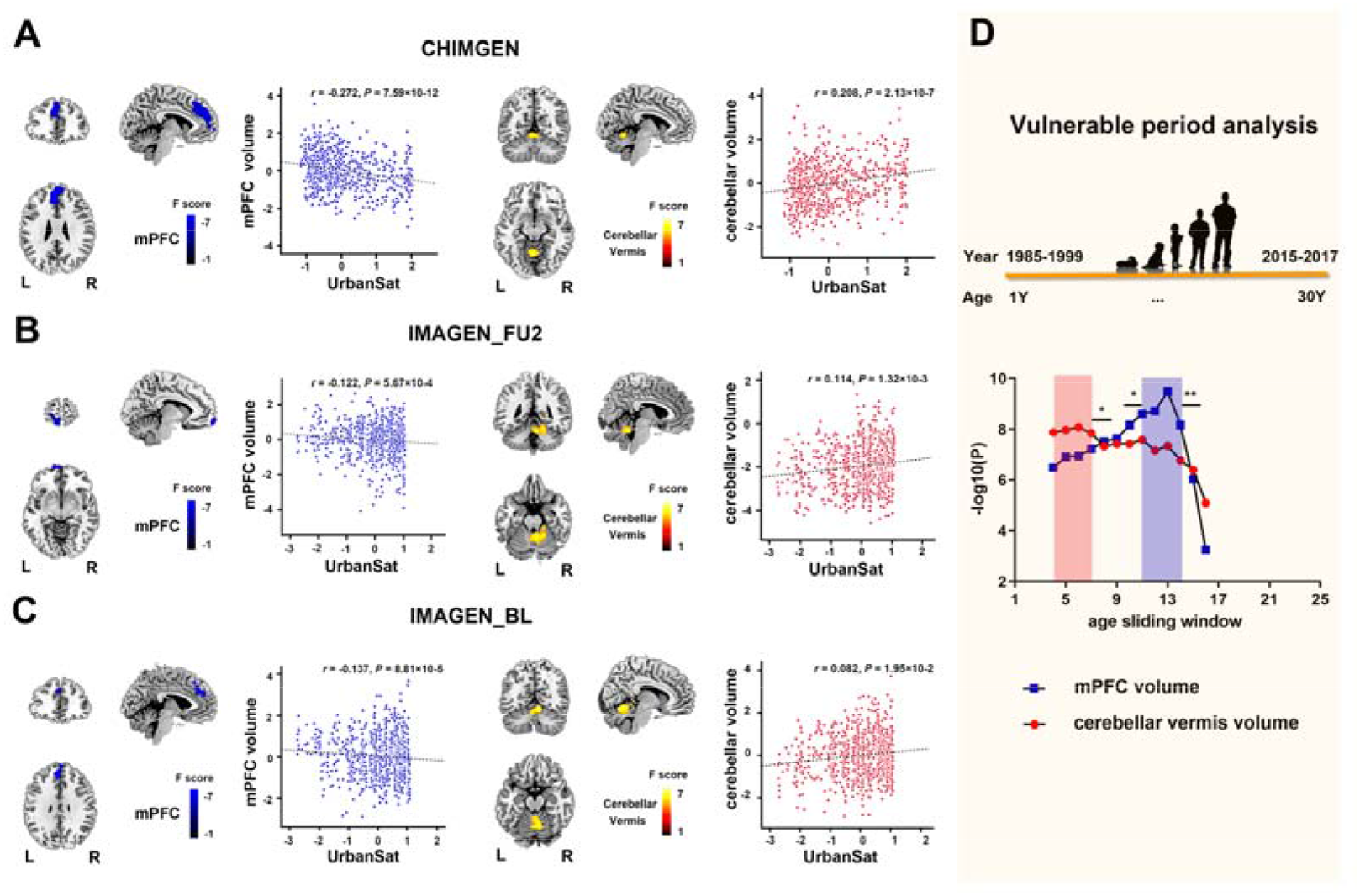
Voxel wise correlation between UrbanSat and GMV across the whole brain. A. In the CHIMGEN sample (N=831), there was a significant negative correlation of the average exposure to urbanicity (measured by UrbanSat before age 18 years) with left mPFC volume (left) and a significant positive correlation with cerebellar vermis volume (right) (FWE-correction, voxel P<0.05); B. In the IMAGEN sample at 19 years(N=791), there was also a significant negative correlation of UrbanSat at 19 years with left mPFC volume (left) and a significant positive correlation with cerebellar vermis volume (right) (FWE-correction, voxel P<0.05); C. The correlations of UrbanSat on brain GMV changes in the left mPFC volume (left) and the cerebellar vermis volume (right) were also present in the IMAGEN baseline sample at 14 years (FWE-correction, voxel P<0.05); D. The correlation of longitudinal UrbanSat with the left mPFC volume was highest during adolescence (ages 11 to 14 years). The difference to the adjacent age bands was significant compared to age 10 years (P=0.012) and to age 15 years (P=2.36×10^-5^); The correlation of longitudinal UrbanSat with cerebellar vermis volume was highest during childhood (ages 4 to 7 years). The difference to the adjacent age bands was significant at P=0.016 compared to age 8 years; BL, baseline datasets at 14 years old; FU2, follow up 2 datasets at 19 years old; L, left; mPFC, medial prefrontal cortex; R, right; * P<0.05; ** P<0.001.

We observed similar results in the ground-level Chinese population grid data^14^, thus validating the effect of urbanicity obtained by satellite measures on brain volume (Supplementary Figure S1). To replicate our findings in a European sample, we carried out a similar voxel-wise multiple regression analysis in 791 adolescents from the IMAGEN at age 19 (19.00 ± 0.71 years) (Supplementary Table S1). UrbanSat also showed a negative correlation of UrbanSat with left mPFC volume (MNI-coordinate: x=-7.5, y=67.5, z=-13.5; 460 voxels; *F*-value peak=-5.71) and a positive correlation with cerebellar vermis volume (MNI-coordinate: x=-5.5, y=-59.5, z=-9.5; 876 voxels; *F* value peak=5.65; Figure 2C) (FWE-correction, voxel *P*<0.05, cluster-size>100 voxels), thus replicating the findings from CHIMGEN. Our findings indicate that the observed correlation between UrbanSat and GMV was independent of geographical locations and socio-cultural conditions.

We then investigated potential susceptibility periods during which exposure to urbanicity may have the strongest influence on brain GMV. In the CHIMGEN young adults, for whom geoposition data were available since birth, we created age sliding windows by averaging UrbanSat for each subject over a period of three years and measured correlations between the exposure to urbanicity in the resulting windows and GMV. The periods showing the greatest susceptibility to urbanicity were childhood (age 4 to 7 years) and adolescence (age 11 and 14 years) (Figure 2B). During childhood, we observed significantly higher correlation of urbanicity with cerebellar vermis volume. While the correlation of longitudinal UrbanSat with the left mPFC volume was highest during adolescence (FWE-correction, voxel *P*<0.05, see Supplementary Results).

While neuroimaging measures in CHIMGEN were acquired once during young adulthood, neuroimaging measures in IMAGEN were acquired twice, during adolescence at 14 years and in a follow-up assessment during early adulthood at 19 years. To further explore susceptibility periods of GMV to the exposure to urbanicity in IMAGEN participants, we analyzed the same individuals at age 14 years (14.05±0.75 years). We confirmed the negative correlation with left mPFC volume (MNI-coordinate: x=-1.5, y=58.5, z=33; 981 voxels; *F* value peak=-5.64) and the positive correlation of UrbanSat with cerebellar vermis volume (MNI-coordinate: x=6, y=-55.5, z=-10.5; 2327 voxels; *F*-value peak=5.04; Figure 2D) (FWE-correction, voxel wise *P*<0.05, cluster-size>100 voxels).

We next investigated the correlation of UrbanSat with white-matter microstructure using tract-based spatial statistics analysis of diffusion tensor imaging (DTI) neuroimaging data. We did not find any significant correlation of UrbanSat with brain fractional anisotropy in either CHIMGEN or IMAGEN (FWE and TFCE-correction, voxel wise *P*<0.05).

### The relation of UrbanSat with functional network connectivity within- and between-brain networks

To investigate the shared and distinct relations of urbanicity with brain activity of cognitive processes in China and Europe, we tested the association of UrbanSat with functional network connectivity (FNC) of 17 within-networks and 136 (17*16/2) between-networks (Figure 3A and Supplementary Methods). In CHIMGEN (N=827), a voxel-wise multiple regression analysis controlling for age, gender and sites revealed a negative correlation of UrbanSat with FNC within the left mPFC of the anterior default mode network (aDMN) (MNI-coordinate: x=6, y=33, z=57; 130 voxels; *F*-value peak=-4.96) and a positive correlation with FNC within the left lingual gyrus of the medial visual network (mVN) (MNI-coordinate: x=-6, y=-84, z=-6; 144 voxels; *F*-value peak=5.21; Figure 3B) (FWE-correction, voxel *P*<0.05, cluster-size>100 voxels). In IMAGEN at 19 years (N=614), UrbanSat showed a negative correlation with FNC within the left mPFC of aDMN (MNI-coordinate: x=-3, y=63, z=6; 106 voxels; *F*-value peak=-4.83; Figure 3D), but not in mVN (FWE-correction, voxel *P*<0.05, cluster-size>100 voxels). The FNC-findings in CHIMGEN and IMAGEN were consistent with the brain GMV changes, indicating that UrbanSat affects both structure and function of the mPFC.

**Figure 3.**
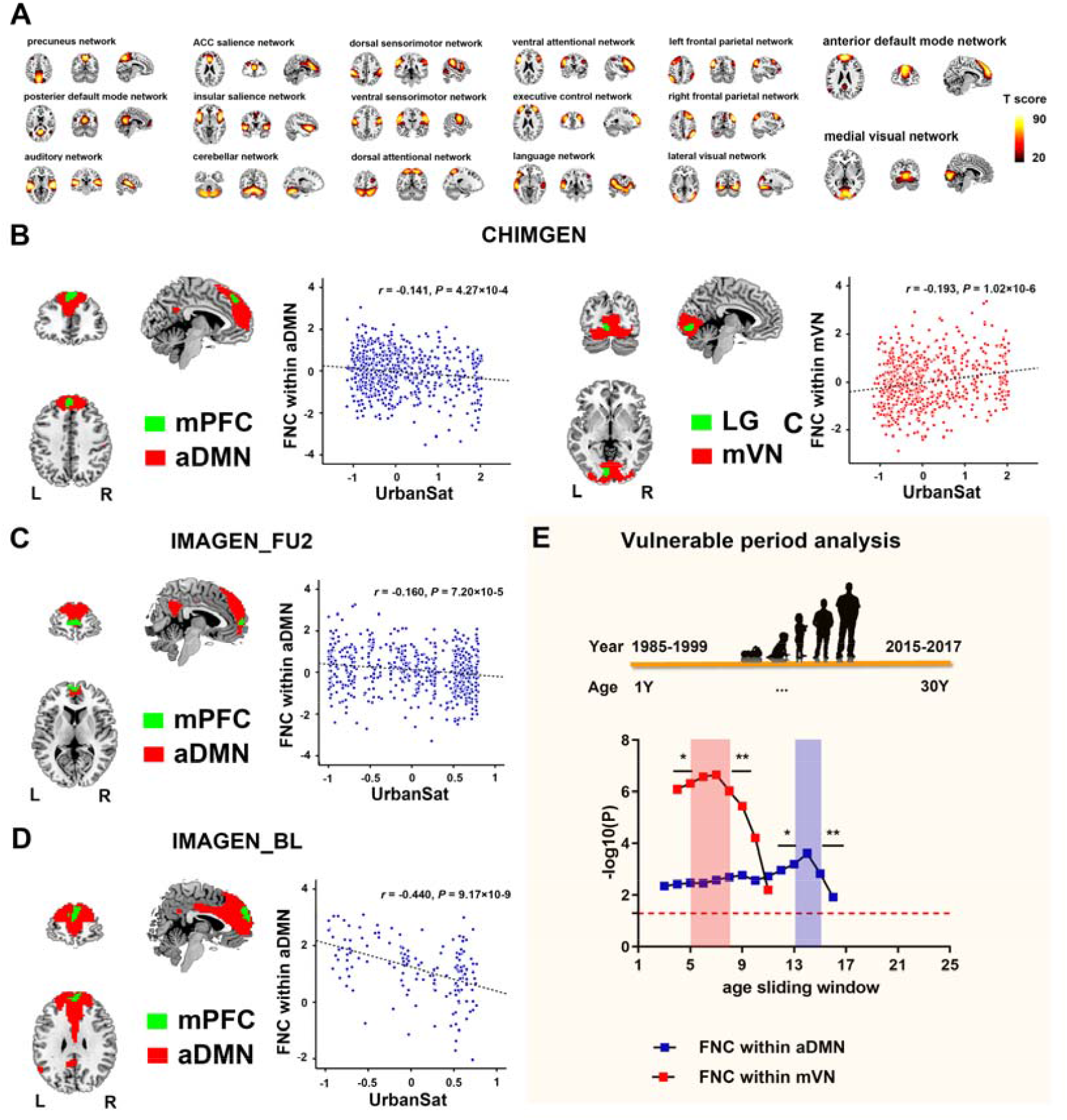
Voxel wise correlations between UrbanSat and FNC within brain networks. A. 17 independent resting state functional networks identified by independent component analysis in the CHIMGEN sample; B. In the CHIMGEN sample (N=827), there was a significant negative correlation of UrbanSat before age 18 years with FNC within the left mFPC of the aDMN (left) and a significant positive correlation with FNC within the left LG of the mVN (right) (FWE-correction, voxel P<0.05); C. In the IMAGEN at age 19 years (N = 614), there was also a significant negative correlation of UrbanSat at 19 years with FNC within the aDMN (FWE-correction, voxel P<0.05) (but not with FNC within the left LG of the mVN); D. The correlation of UrbanSat on FNC within the aDMN was also present in the IMAGEN at 14 years (FWE-correction, voxel P<0.05). E. The correlation of UrbanSat with FNC within the aDMN was highest during adolescence (ages 13 to 15 years). The difference to the adjacent age bands was significant compared to age 12 years (P=0.010) and to age 16 years (P=6.16×10^-5^); The correlation of UrbanSat with FNC within the mVN was only vulnerable in childhood and highest during childhood (ages 5 to 8 years). The difference to the adjacent age bands was significant at P=0.027 compared to 4 years and P=2.68×10^-4^ compared to age 9 years; aDMN, anterior default mode network; FNC, functional network connectivity; L, left; LG, lingual gyrus; mPFC, medial prefrontal cortex; mVN, medial visual network; R, right; * P<0.05; ** P<0.001.

Upon investigating potential susceptibility periods for FNC within the aDMN, we found the greatest correlation with UrbanSat during adolescence (age 13 to 15 years) (FWE-correction, voxel *P*<0.05, Figure 3C and see Supplementary Results). We confirmed the relation of UrbanSat with the FNC of the left mPFC within the aDMN in the IMAGEN participants at age 14 years (MNI-coordinate: x=0, y=60, z=24; 104 voxels; *F* value peak=-5.61; Figure 3E) (FWE-correction, voxel *P*<0.05, cluster-size>100 voxels). Together, our data suggest a susceptibility window for the correlation of urbanicity and FNC within aDMN in adolescence. In the case of the mVN, we found the greatest correlation with UrbanSat from age 5 to 8 years, indicating a susceptibility window in childhood (Figure 3C).

In the resulting 136 between-network FNCs we found overall similarity of resting-state activity between CHIMGEN and IMAGEN (Supplementary Figure S2). For the 136 pairs of between-network FNCs, However, UrbanSat was significantly correlated with 42 connections in CHIMGEN, 19 connections in IMAGEN at age 19 and 27 connections in the same IMAGEN participants at age 14 (Figure 4A-C) (*P*<0.05; to account for correlations in between-network connectivity we carried out 10000 permutations), suggesting a greater sensitivity to urbanicity in the Chinese CHIMGEN sample compared to the European IMAGEN sample. Five connections from six brain networks (aDMN, anterior default mode network; CBN, cerebellar network; ECN, executive control network; LN, language network; rFPN/lFPN, right or left frontal parietal network) were shared between CHIMGEN and IMAGEN: aDMN-CBN, aDMN-rFPN, aDMN-ECN, rFPN-LN and rFPN-lFPN (Figure 4D and Supplementary Table S2 and Figure S3). Whereas the susceptibility period for between-network connectivity involving self-referential thoughts^22^ and executive control^23^, such as aDMN-CBN, aDMN-rFPN, aDMN-ECN, and lFPN-rFPN was during adolescence (Figure 4E and Supplementary Results), between-network connectivity involving the language network LN-rFPN was most susceptible to urbanicity during childhood (Figure 4E and Supplementary Resutls). These results indicate that between-networks FNCs relate to urbanicity in both shared and distinct ways during brain development and in different socio-cultural and geographic locations.

**Figure 4.**
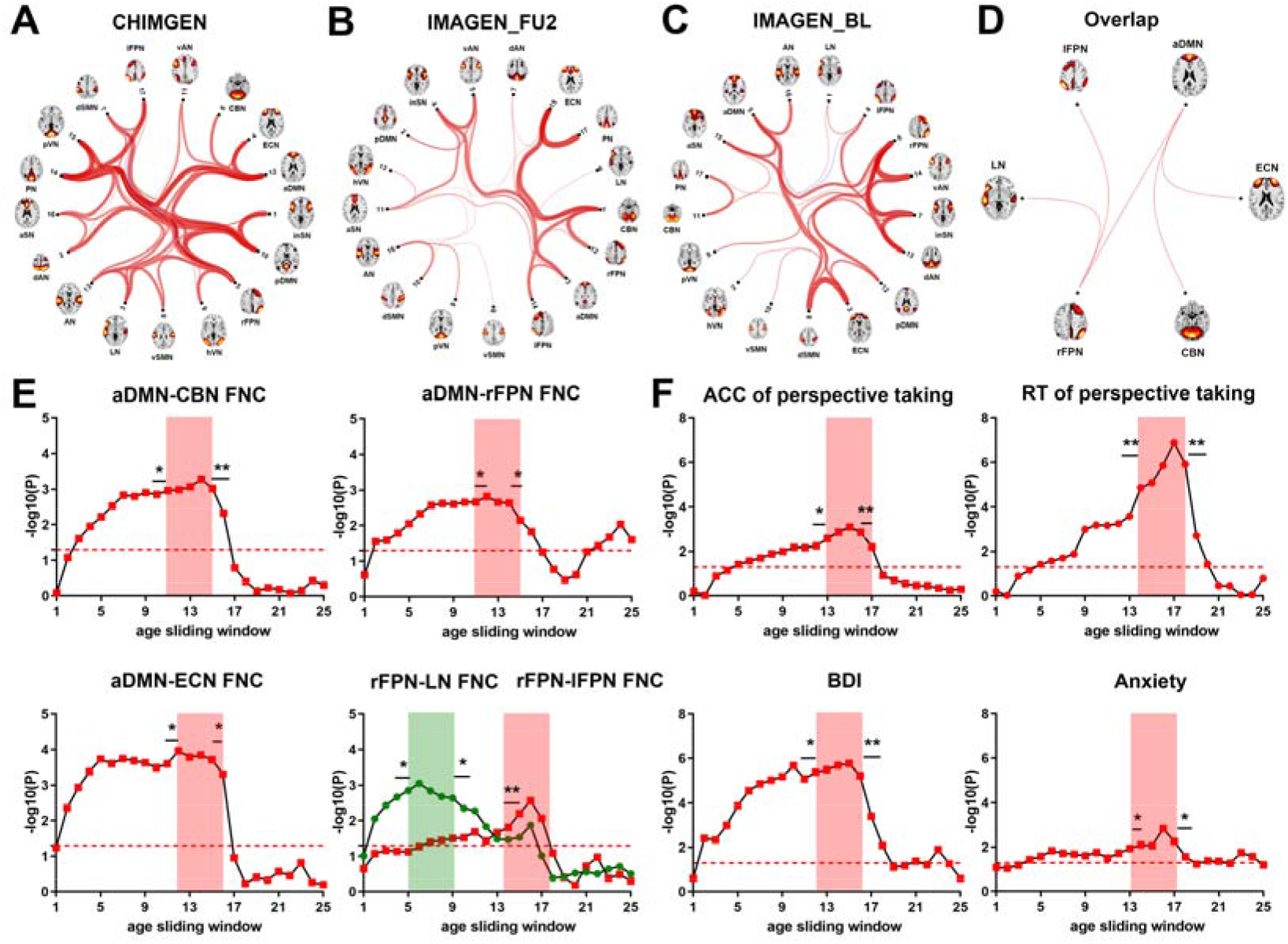
Correlations of UrbanSat with between-networks FNCs and behaviors. A. In the CHIMGEN study, UrbanSat was significantly correlated with 42 brain between-networks FNCs; B. In the IMAGEN FU2 study at age 19 years, UrbanSat was significantly correlated with 19 brain between-networks FNCs; C. In the IMAGEN BL study at age 14 years, UrbanSat was significantly correlated with 27 brain between-networks FNCs; D. Only five pairs of brain between-networks FNCs could be replicated in CHIMGEN and IMAGEN sample; E. The correlation of longitudinal UrbanSat on five overlapping brain between-networks FNCs was significantly increased in the childhood period (age 5 to 9 years in rFPN with LN) and adolescent period (age 11 to 15 years in aDMN with CBN, age 12 to 14 years in aDMN with rFPN, age 12 to 15 years in aDMN with ECN, and age 15 to 17 years in rFPN with lFPN) in the CHIMGEN sample; F. In the CHIMGEN sample, the correlation of longitudinal UrbanSat with perspective taking performance was significantly increased in the adolescent period (age 13 to 16 years in accuracy and age 14 to 18 years in reaction time). The difference to the adjacent age bands was significant in accuracy (P=0.026 compared to age 12 years and P=1.45×10^-4^ compared to age 17 years) and in reaction time (P=6.58×10^-5^ compared to age 13 years and P=1.16×10^-6^ compared to age 19 years); The correlation of longitudinal UrbanSat on mental health was significantly increased in the adolescent period (age 12 to 16 years in depressive severity and age 14 to 17 years in state anxiety). The difference to the adjacent age bands was significant in BDI (P=0.016 compared to age 11 years and P=3.62×10^-5^ compared to age 17 years) and state anxiety (P=0.036 compared to age 13 years and P=0.011 compared to age 18 years). ACC, accuracy; aDMN, anterior default mode network; BDI, Beck Depression Inventory; CBN, cerebellar network; ECN, executive control network; FNC, functional connectivity; LN, language network; lFPN, left frontal parietal network; rFPN, right frontal parietal network; RSQ, Ruminating Scale Questionnaire; RT, reaction time; * P<0.05; ** P<0.001.

### The relation of UrbanSat with behavior

We investigated whether UrbanSat was correlated with measures of cognition and mental health-relevant behavior, in particular symptoms of depression and anxiety. Of the domains of neuropsychology, only one measure of social cognition, namely perspective taking, a measure of perceiving a situation or understanding a concept from an alternative point of view^24^, was significantly positively associated with UrbanSat (accuracy: *r*=0.124, *P=*0.002; reaction time, *r*=-0.245, *P*= 6.10×10^-7^; Table 1 and Supplementary Figure S4) (Bonferroni-correction *P*<0.05). The correlation between UrbanSat and perspective taking performance was strongest during adolescence (age 13 to 16 years in accuracy and 14 to 18 years in reaction time). We confirmed the positive correlations between UrbanSat and perspective taking in IMAGEN at age 16 years (*r*=0.103, *P=*0.009), the earliest age these data were available (Figure 4F, Table 2 and Supplementary Figure S4). In the IMAGEN at age 19 years, we observed a trend-level significant correlation between UrbanSat and perspective taking (*P*=0.056) (Table 2 and Supplementary Figure S4).

**Table 1.**
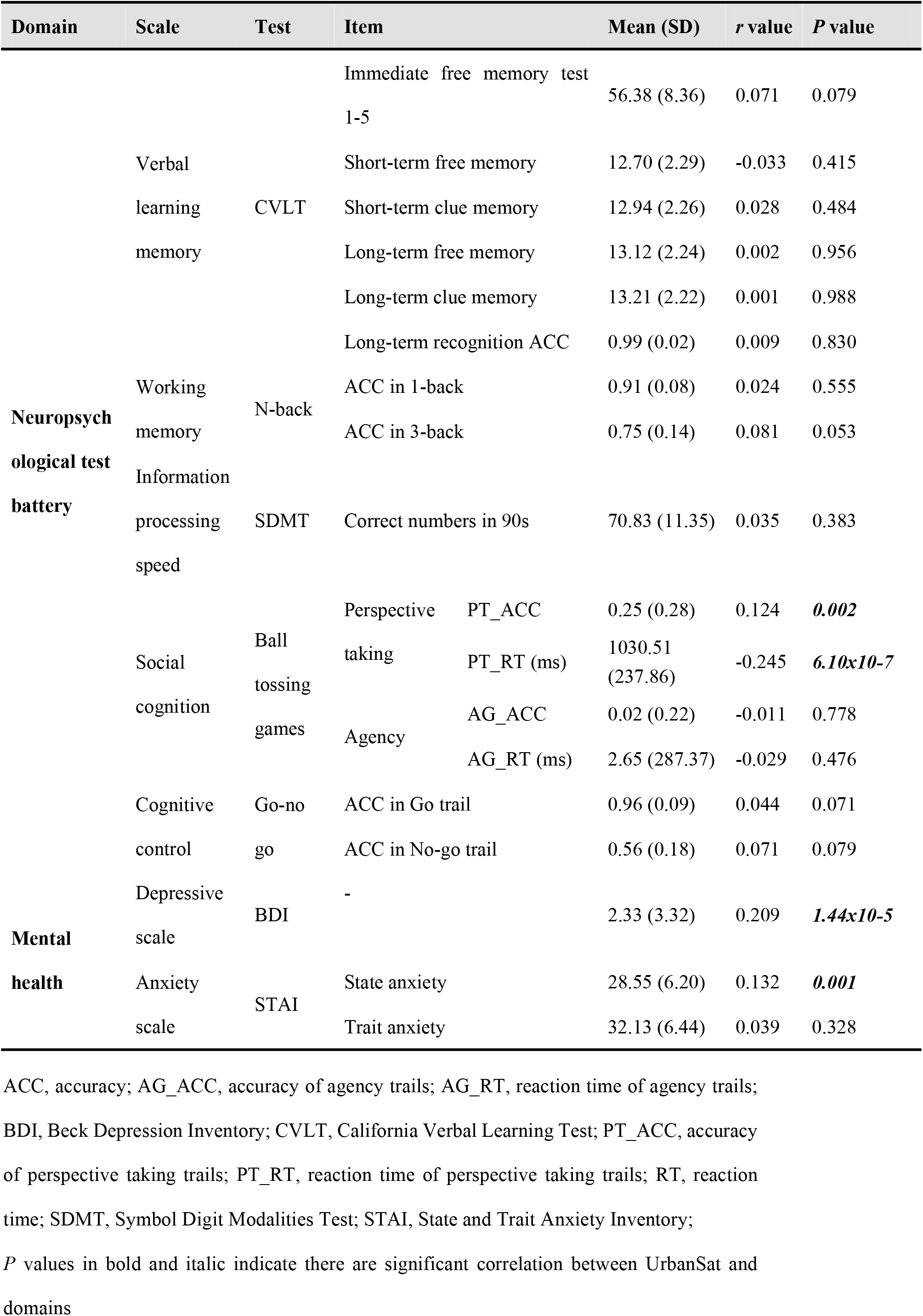
The correlation of UrbanSat with neuropsychological assessments and mental health in CHIMGEN sample (N = 622)

**Table 2.** The correlation of UrbanSat with IRI and mental health in IMAGEN sample.

| Scale | Item | N | Mean (SD) | *r* value | *P* value |
| --- | --- | --- | --- | --- | --- |
| IRI FU1 | Empathic concern | 636 | 16.17 (3.84) | 0.022 | 0.574 |
|  | Fantasy | 636 | 17.08 (4.67) | 0.030 | 0.457 |
|  | Perspective taking | 636 | 17.96 (3.78) | 0.103 | **0.009** |
|  | Personal distress | 636 | 12.06 (3.81) | -0.036 | 0.372 |
| IRI FU2 | Empathic concern | 572 | 17.12 (4.67) | 0.067 | 0.109 |
|  | Fantasy | 572 | 17.12 (3.79) | -0.01 | 0.819 |
|  | Perspective taking | 572 | 18.59 (3.90) | 0.08 | 0.056 |
|  | Personal distress | 572 | 11.38 (3.82) | 0.049 | 0.243 |
| Depressive scale FU2 | RSQ | 575 | 37.90 (12.03) | 0.118 | **0.004** |
| Anxiety scale FU2 | DAWBA-GA | 842 | 1.43 (0.76) | 0.059 | 0.122 |
|  | CIDI-AS | 813 | 1.26 (0.63) | 0.035 | 0.320 |

CIDI-AS, Anxiety Screening from the Composite International Diagnostic Interview; DAWBA-GA, Generalized Anxiety Scale from The Development and Well-Being Assessment Interview; FU1, IMAGEN follow up 1 assessment acquired at 16 years; FU2, IMAGEN follow up 2 assessment acquired at 19 years; N, sample size; IRI, Interpersonal Reactivity Index; RSQ, Ruminating Scale Questionnaire; SD, stand deviation.

In CHIMGEN, UrbanSat was correlated with number of symptoms in the Beck Depressive Inventory (BDI) (*r* = 0.209, *P=*1.44×10^-5^) and state anxiety (*r* = 0.132, *P=*0.001) (Bonferroni *P* < 0.05) (Table 1 and Supplementary Figure S4). The correlations between UrbanSat with BDI and state anxiety suggested greatest susceptibility during adolescence. (Figure 4F and Supplementary Results). In IMAGEN, we validated the correlation between UrbanSat and depressive symptoms at age 19 (*r*=0.118, *P=*0.004) using the Ruminating Scale Questionnaire (RSQ) (No depression questionnaire was available at age 14 or 16). There was no association between UrbanSat and anxiety at any timepoint in IMAGEN (Table 2 and Supplementary Figure S4).

### Mediation of the correlation of UrbanSat with behavior by brain measures

We then investigated if left mPFC and cerebellar vermis volume, as well as the significant within- and between-network connectivity mediate the correlation of UrbanSat with perspective taking and symptoms of depression and anxiety.

In CHIMGEN, the correlation of UrbanSat with perspective taking was mediated by the left mPFC volume (*P<*0.001, *r*^2^=0.20), cerebellar vermis volume (*P<0.001, r*^2^=0.15), FNC within aDMN (*P*<0.001, *r*^2^=0.12) and VN (*P*<0.001, *r*^2^=0.13), as well as by the between network FNCs aDMN-CBN (*P<0.05*, *r*^2^=0.10), aDMN-rFPN (*P<0.001*, *r*^2^=0.13) (Figure 5A). In IMAGEN at age 16, the association of UrbanSat with perspective taking was also mediated by left mPFC volume (*P*<0.05, *r*^2^=0.10), FNC within the aDMN (*P*<0.001, *r*^2^=0.36), between network FNC aDMN-CBN (*P<0.001*, *r*^2^=0.23), aDMN-rFPN (*P<0.001*, *r*^2^=0.20) (Figure 5D). We did not find a mediation effect of the cerebellar vermis volume in the IMAGEN at age 16 (*P=*0.26).

**Figure 5.**
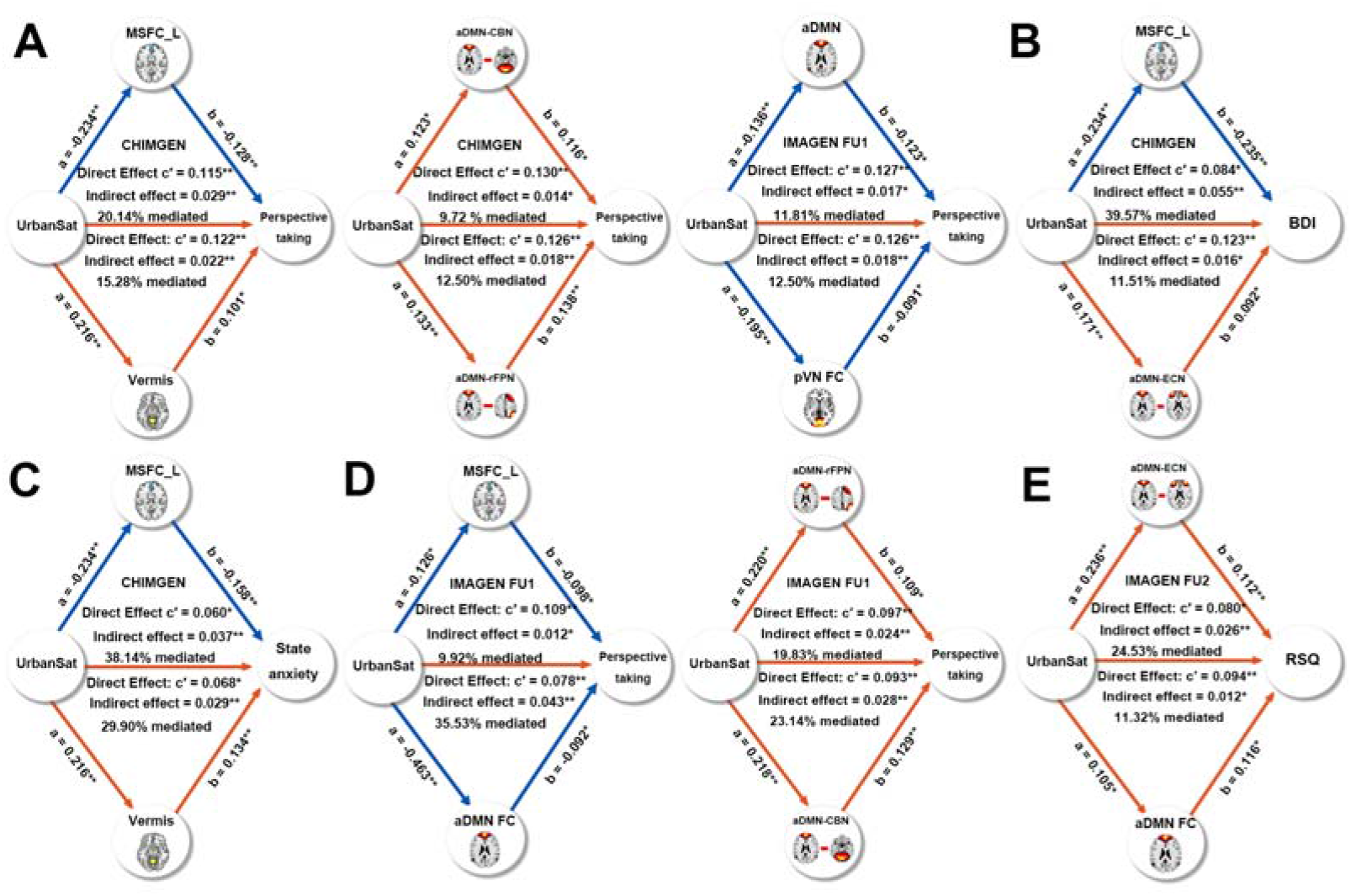
Mediation analysis in the UrbanSat-brain-behavious pathway. A. In the CHIMGEN sample, the left mPFC and cerebellar vermis volume, FNC within the aDMN and the VN, as well as between-networks FNC of the aDMN and CBN and the aDMN and rFPN mediated the effect of UrbanSat on perspective taking performance; B. The left MPFC volume and between-networks FNC of the aDMN with ECN mediated the effect of UrbanSat on BDI; C. The left mPFC and cerebellar vermis volume mediated the effect of UrbanSat on state anxiety; D. In the IMAGEN FU1 assessment at 16 years, but not in the FU2 at 19 years, the effect of UrbanSat on perspective taking was mediated by the left MPFC volume, FNC within the aDMN, between-networks FNC of the aDMN with CBN and the aDMN with rFPN; E. In the IMAGEN FU2 assessment at 19 years, the effect of UrbanSat on RSQ was mediated by FNC within the aDMN, between-networks FNC of the aDMN with ECN. RSQ data were not available at IMAGEN BL sample at 14 years. aDMN, anterior default mode network; CBN, cerebellar network; ECN, executive control network; FNC, functional connectivity; FPN, frontal parietal network; BDI, Beck Depression Inventory; RSQ, Ruminating Scale Questionnaire; * P<0.05; ** P<0.001.

In CHIMGEN, the correlation between UrbanSat and BDI was mediated by left mPFC volume (*P<0.001*, *r*^2^=0.40) and aDMN-ECN (*P<0.05*, *r*^2^=0.12) but not by the cerebellar vermis volume (*P=*0.97) (Figure 5B). Susceptibility to state anxiety was mediated by the left medial frontal superior cortex volume (*P<0.001*, *r*^2^=0.38) as well as by cerebellar vermis volume (*P<0.001*, *r*^2^=0.30) (Figure 5C). In IMAGEN the correlation between UrbanSat and depressive symptoms was mediated by FNC within the aDMN (*P<0.05*, *r*^2^=0.11) and aDMN-ECN (*P<0.001*, *r*^2^=0.25) (Figure 5E), but not by the left mPFC (*P=*0.09) or by the cerebellar vermis (*P=*0.16).

## Discussion

Using innovative remote-sensing satellite data acquisition, we provide a comparative characterization of the relation of urbanicity, brain structure and function, cognition and behavior in large datasets of young people in China and Europe. We found shared correlations of urbanicity with brain structure and function, as well as perspective taking, a measure of social cognition, and symptoms of depression in Chinese CHIMGEN and European IMAGEN sample. In contrast, correlations with anxiety were evident only in the Chinese sample. Brain structure analyses revealed a negative correlation of urbanicity with volume of the mPFC, with peaks in the dorsal (dmPFC) and ventral part (vmPFC) in Chinese and Europeans, respectively. This correlation was significantly greater in adolescence, with strongest correlations at age 13 years. The mPFC mediated the effect of UrbanSat on perspective taking in CHIMGEN and IMAGEN, and the effect on depression and anxiety in CHIMGEN, only. Reduction of mPFC structure is implicated in responses to chronic stress^25^ and pollution^26^, both associated with urban living. The areas within the mPFC showing the greatest correlation with urbanicity in CHIMGEN and IMAGEN, the dmPFC and vmPFC, are thought to be involved in social and affective behavior^27^.

We also found a positive correlation of urbanicity with cerebellar vermis volume in both Chinese and Europeans, with the strongest correlation during childhood at age 6. Lesions in the vermis give rise to the ‘Cerebellar-Cognitive-Affective-Syndrome’, characterized by impairments in executive function and memory, as well as affective behavior^28^. Animal studies extend these findings to impairments in social behavior^29^ and stress-dependent depressive affect^30^, involving dopaminergic^29^ and serotonergic^30^ mechanisms, thus providing candidate neurotransmitter systems for urbanicity investigations. Vermis volume mediates the effect of urbanicity on perspective taking and anxiety in Chinese only, suggesting distinct structural brain characteristics mediating urbanicity in China and Europe. Together, our findings relate urbanicity to structural brain changes in regions linked to shared and distinct social and affective behavior among individuals from different sociocultural conditions and geographical locations. Our results are supported by reports of associations of urbanicity and prefrontal cortex, in particular mPFC^31^ and DLPFC^32^, in smaller European studies.

We used FNC of 17 resting-state networks related to cognitive and sensory-motor processes to measure shared and distinct network connectivity in Chinese and Europeans. The correlation of urbanicity with FNC within the aDMN was shared between CHIMGEN and IMAGEN. FNC within the aDMN mediated the effect of UrbanSat on perspective taking, both in CHIMGEN and IMAGEN, and on depressive symptoms in IMAGEN only. Similarly to mPFC, FNC within the aDMN has been involved in social cognition^33^ and depression^34^, and is also moderated by urbanicity-related risk factors, including chronic stress^35^ and air pollution^36^. FNC within the mVN, which might be involved in the organization of cognitive function^37^, was associated with urbanicity in CHIMGEN only, with a pronounced peak in susceptibility at the age of 6 years. It also mediated the effect of urbanicity on perspective taking, again in CHIMGEN only.

The resulting 136 between-network FNCs demonstrated overall similarity of resting-state activity between CHIMGEN and IMAGEN (Supplementary Figure S2). Of note, urbanicity was specifically correlated with 42 connections in CHIMGEN, 19 connections in IMAGEN at age 19 and 27 connections in IMAGEN at age 14. Of the 5 between-network FNCs correlated with urbanicity present in both CHIMGEN and IMAGEN, aDMN-ECN FNC mediated the effect of UrbanSat on depressive symptoms in both samples. Consistent with the mPFC structural and within FNC changes in the aDMN, FNC between aDMN and ECN have been implicated in the emotional dysregulation of depression^38^. aDMN-rFPN FNC and aDMN-CBN FNC, shown to be engaged in social cognition^39, 40^, mediated the effect of UrbanSat on perspective taking in both CHIMGEN and IMAGEN. It is possible that a combination of structural changes in the mPFC and cerebellum account for this altered functional connectivity. Together, they may mediate the relation of urbanicity with perspective taking. The majority of between FNC associated with urbanicity was distinct between CHIMGEN and IMAGEN at age 19 and 14, however, indicating a specificity of between-network connectivity across geographic locations as well as brain developmental stage. Furthermore, between-network connectivity was increased in the Chinese sample, suggesting a greater sensitivity to urbanicity. This might reflect the more drastic changes in urbanization in China compared to Europe^41^, but may also relate to ethnically distinct genetic and environmental interactions, involving different brain pathways^42^.

Whereas previous studies have investigated urban living before the age of 15 years^43^, our characterization of specific age windows of increased susceptibility indicate discrete neurodevelopmental periods during which brain development is sensitive to the effect of urbanicity. The susceptibility period for the relation of urbanicity with mPFC-volume during adolescence coincides with increased synaptic pruning, which depends (in part) on environmental stimulation of neurons^44^. It is conceivable that adverse social situations that elevate stress in an urban environment might result in accelerated synaptic pruning^45^, which might also affect functional network connectivity. Among the shared between-network connections, those that involve the aDMN and mediate perspective taking and depression show increased susceptibility for urbanicity during adolescence. The susceptibility period of vermis around age 6 years is consistent with the high volume increase of midline cerebellar structures observed in this age group^46^. The fact that behavioral manifestations mediated by cerebellar vermis are all associated with susceptibility windows during adolescence may indicate an early neurobehavioral trajectory with delayed behavioral manifestation^47^.

In contrast to previous studies reporting an association of urbanicity with social and cognitive impairments^48^, we find better performance of one social cognition variable, perspective taking, in an urban environment. These previous findings were mostly derived from samples at risk for schizophrenia^48^. As our cohorts were recruited from the general population, it is possible that we detect an adaptive behavior to living in the city, which might be compromised in patients at risk for psychosis, thus exacerbating their social difficulties and isolation in an urban environment^49^. We are extending previous observations reporting an increased incidence of depression and anxiety symptoms in urban settings^50^, by demonstrating the stability of this observation in different geographical and sociocultural regions, and by discovering possible underlying brain mechanisms.

Remote sensing satellites play a critical role in monitoring the Earth’s surface and atmosphere to track environmental conditions that are intimately related to human health^51^. Satellites have been applied to map urbanization, poverty, climate change and pollution^52^, as well as the spread of infectious disease^53^. By extending this approach to measuring the relation of urbanicity with brain development, cognition and mental health, our study provides an approach that might be useful to characterize and monitor the spatial and temporal patterns of risk for mental disorders. Advances in resolution of satellite and computing power of geographic system facilitate integration of remote-sensing environmental parameters with public health data to develop models for disease surveillance and control. These models might be useful for sustainable urban planning and policy, and to develop targeted prevention efforts for young people in their unique environment. Our satellite measure of population density does not capture with sufficient granularity important social environmental conditions resulting from increased population density, such as social stress^54^, access to infrastructure and green space^55^ or crime^56^. Future studies will investigate the integrated effect of urban physical and social environment, as well as the interaction with genetics and their relation to brain and behavior. As our approach can be extended and generalized to other geographic locations and is easy to implement even in the absence of detailed or directly comparable ground level data, it may be relevant for public health, policy and urban planning globally.

## Acknowledgments

This work was partly supported by the National Key Research and Development Program of China (Grant No. 2018YFC1314301 and Grant No. 2017YFA0604401), the National Natural Science Foundation of China (Grant No. 81425013), the European Union-funded FP6 Integrated Project IMAGEN (Reinforcement-related behaviour in normal brain function and psychopathology) (LSHM-CT-2007-037286), the Horizon 2020 funded ERC Advanced Grant ‘STRATIFY’ (Brain network based stratification of reinforcement-related disorders) (695313), ERANID (Understanding the Interplay between Cultural, Biological and Subjective Factors in Drug Use Pathways) (PR-ST-0416-10004), BRIDGET (JPND: BRain Imaging, cognition Dementia and next generation GEnomics) (MR/N027558/1), the FP7 project MATRICS (603016), the Medical Research Council Grant ‘c-VEDA’ (Consortium on Vulnerability to Externalizing Disorders and Addictions) (MR/N000390/1), the National Institute for Health Research (NIHR) Biomedical Research Centre at South London and Maudsley NHS Foundation Trust and King’s College London, the Bundesministeriumfür Bildung und Forschung (BMBF grants 01GS08152; 01EV0711; eMED SysAlc01ZX1311A; Forschungsnetz AERIAL 01EE1406A, 01EE1406B), the Deutsche Forschungsgemeinschaft (DFG grants SM 80/7-2, SFB 940/2), the Medical Research Foundation and Medical research council (grant MR/R00465X/1), the Human Brain Project (HBP SGA2). Further support was provided by grants from: ANR (project AF12-NEUR0008-01 - WM2NA, and ANR-12-SAMA-0004), the Fondation de France, the Fondation pour la Recherche Médicale, the Mission Interministérielle de Lutte-contre-les-Drogues-et-les-Conduites-Addictives (MILDECA), the Assistance-Publique-Hôpitaux-de-Paris and INSERM (interface grant), Paris Sud University IDEX 2012, the National Institutes of Health, Science Foundation Ireland (16/ERCD/3797), U.S.A. (Axon, Testosterone and Mental Health during Adolescence; RO1 MH085772-01A1), NIH grant R01EB020407 and R01EB006841 and by NIH Consortium grant U54 EB020403, supported by a cross-NIH alliance that funds Big Data to Knowledge Centres of Excellence.

## Methods

### CHIMGEN and IMAGEN project

In Chinese Imaging Genetics project (CHIMGEN), 7200 healthy Chinese Han participants of 18-30 ages with blood sample, environmental, structural and functional neuroimaging and cognitive assessment were recruited from 29 centers of Chinese mainland 20 cities. In IMAGEN study, the first European multisite and longitudinal project^13^, comprehensive environmental factor, genetics, transcriptome, epigenetics, structural and functional neuroimaging, neurocognitive measure and mental health outcome were collected from more than 2000 14-year-old adolescents and 19-year-old follow up young adults.

### Remote sensing satellite data

Global Human Settlement Layer (GHSL)^14^, Nighttime Lights (NL)^15^, Normalized Difference Vegetation Index (NDVI)^16^, Normalized Difference Built-up Index (NDBI)^17^, Normalized Difference Water Index (NDWI)^18^ and global land cover mapping^19^ were extracted from Google Earth Engine to measure different urbanicity characteristics based on the acquired individual geographic position from CHIMGEN (n=3306) and IMAGEN (n=1205) (Supplementary Methods and Table S3-S5).

### Neuroimaging data

In this study, 1104 subjects from CHIMGEN Tianjin main centers, and 1724 and 1423 subjects from IMAGEN (at 14 years and 19 years, respectively) with available neuroimaging data were included in the further analysis. In both studies, T1-weighted imaging, DTI and resting-state functional imaging were acquired using 3.0 Tesla MRI scanners. Brain GMV from the T1-weighted scans, white matter tract based spatial statistics (TBSS) from the DTI and within and between functional network connectivity (FNC) from the resting-state functional imaging were calculated from preprocessed neuroimaging data (Supplementary Methods and Table S1). Only subjects where both quality-controlled neuroimaging data and remote sensing satellite data were available were included in the statistical analysis.

### Neuropsychological assessment and mental health data

Verbal learning memory, working memory, information processing speed, social cognition and executive control were included in the neuropsychological assessment, and depression and anxiety were included in mental health, in both CHIMGEN and IMAGEN (Supplementary Methods and Figure S5).

### Statistical analysis

To test the latent urbanicity variable derived from the satellite features, confirmatory factor analysis (CFA) was applied to the measuring satellite features using R package *lavaan*^20^. Only the closely correlated satellite features (*r* > 0.4) were included in the CFA model used to construct a factor for urbanicity in China and Europe respectively, which we termed ‘UrbanSat’ (Supplementary Table S6). Population density is the measure currently typically used to map urbanicity processes in sociology^21^, so to consolidate our foundation of using satellite features to measure urbanicity, the UrbanSat was correlated with this “gold standard” ground-level population grid GHSL data from both China and Europe, separately (Supplementary Figure S6). The detailed statistical analysis of the relation between UrbanSat and brain GMV, fractional anisotropy (FA), within- and between- FNC of brain networks are described in the Supplementary Methods.

