## Supplementary Material for "Satellite Imaging of Global Urbanicity relate to Adolescent Brain Development and Behavior"

210008 Nanjing, China

[11] Department of Radiology, Xiangya Hospital, Central South University, 410008 Changsha, China

[12] Department of Medical Imaging, The First Affiliated Hospital of Guangzhou University of Chinese Medicine, 510405 Guangzhou, China

[13] Department of Radiology, The First Hospital of Shanxi Medical University, 030001 Taiyuan, China

[14] Department of Radiology, The Second Affiliated Hospital of Zhejiang University, School of Medicine, 310009 Hangzhou, China

[15] Department of Radiology, The First Affiliated Hospital of Anhui Medical University, 230022 Hefei, China

[16] Department of Radiology, Yantai Yuhuangding Hospital, 264000 Yantai, China

[17] Department of Radiology, Tianjin Huanhu Hospital, 300350 Tianjin, China

[18] Functional and Molecular Imaging Key Lab of Shaanxi Province & Department of Radiology, Tangdu Hospital, the Military Medical University of PLA Airforce (Fourth Military Medical University), 710038 Xi'an, China

[19] Department of Radiology, Hainan General Hospital, 570311 Haikou, China

[20] Department of Radiology, Beijing Tongren Hospital, Capital Medical University, 100730 Beijing, China

[21] Department of Radiology, The First Affiliated Hospital of Wenzhou Medical University, 325000 Wenzhou, China

[22] Department of Magnetic Resonance, Lanzhou University Second Hospital, 730050 Lanzhou, China

[23] Department of Psychology, University of Chinese Academy of Sciences (CAS), 100049 Beijing, China

[24] Department of Radiology, Qilu Hospital of Shandong University, 250012 Jinan, China

[25] Department of Radiology, Tianjin First Center Hospital, 300192 Tianjin, China

[26] Department of Radiology, The First Affiliated Hospital of Dalian Medical University, 116011 Dalian, China

[27] Department of Radiology, Pingjin Hospital, Logistics University of Chinese People's Armed Police Forces, 300162 Tianjin, China

[28] Department of Radiology, the Center for Medical Imaging, West China Hospital of Sichuan University, 610041 Chengdu, China

[29] School of Life Sciences, University of Science & Technology of China, 230026 Hefei, China

[30] Department of Radiology, The Affiliated Hospital of Xuzhou Medical University, 221006 Xuzhou, China

[31] Department of Medical Imaging, Jinling Hospital, Medical School of Nanjing University,210002 Nanjing, China
[32] Department of Radiology, Tianjin Medical University Cancer Institute and Hospital, 300060 Tianjin, China
[33] Department of Child and Adolescent Psychiatry and Psychotherapy, Central Institute of Mental Health, Medical Faculty Mannheim, Heidelberg University, Square J5, 68159 Mannheim, Germany;
[34] Department of Neuroimaging, Institute of Psychiatry, Psychology & Neuroscience, King's College London, United Kingdom;
[35] Discipline of Psychiatry, School of Medicine and Trinity College Institute of Neuroscience, Trinity College Dublin, Dublin, Ireland;
[36] Department of Cognitive and Clinical Neuroscience, Central Institute of Mental Health, Medical Faculty Mannheim, Heidelberg University, Square J5, Mannheim, Germany;
[37] Department of Psychology, School of Social Sciences, University of Mannheim, 68131 Mannheim, Germany;
[38] NeuroSpin, CEA, Université Paris-Saclay, F-91191 Gif-sur-Yvette, France;
[39] Departments of Psychiatry and Psychology, University of Vermont, 05405 Burlington, Vermont, USA;
[40] Sir Peter Mansfield Imaging Centre School of Physics and Astronomy, University of Nottingham, University Park, Nottingham, United Kingdom;
[41] Charité – Universitätsmedizin Berlin, corporate member of Freie Universität Berlin, Humboldt-Universität zu Berlin, and Berlin Institute of Health, Department of Psychiatry and Psychotherapy, Campus Charité Mitte, Charitéplatz 1, Berlin, Germany
[42] Physikalisch-Technische Bundesanstalt (PTB), Abbestr. 2 - 12, Berlin, Germany;
[43] Institut National de la Santé et de la Recherche Médicale, INSERM Unit 1000 "Neuroimaging & Psychiatry", University Paris Sud, University Paris Descartes - Sorbonne Paris Cité; and Maison de Solenn, Paris, France;
[44] Institut National de la Santé et de la Recherche Médicale, INSERM Unit 1000 "Neuroimaging & Psychiatry", University Paris Sud, University Paris Descartes - Sorbonne Paris Cité; and Psychiatry Department 91G16, Orsay Hospital, France;
[45] Institut National de la Santé et de la Recherche Médicale, UMR 992 INSERM, CEA, Faculté de médecine, Université Paris-Sud, Université Paris-Saclay, NeuroSpin, F-91191 Gif-sur-Yvette, France;
[46] Bloorview Research Institute, Holland Bloorview Kids Rehabilitation Hospital and Departments of Psychology and Psychiatry, University of Toronto, Toronto, Ontario, M6A 2E1, Canada;
[47] Department of Child and Adolescent Psychiatry and Psychotherapy, University Medical

Centre Göttingen, von-Siebold-Str. 5, 37075, Göttingen, Germany;
[48] Department of Psychiatry and Neuroimaging Center, Technische Universität Dresden, Dresden, Germany;
[49] School of Psychology and Global Brain Health Institute, Trinity College Dublin, Ireland;
[50] School of Global Policy and Strategy, International Lane, University of California San Diego, San Diego, CA 92093;
[51] Center for Wireless and Population Health Systems, Department of Family and Preventive Medicine and Calit2's Qualcomm Institute, University of California, San Diego, 9500 Gilman Drive, Dept. 0811, La Jolla, CA 92093-0811;
[52] Tri-institutional Center for Translational Research in Neuroimaging and Data Science (TReNDS) [Georgia State University, Georgia Institute of Technology, Emory University], Atlanta, GA 30303
[53] Collaborative Innovation Center of Tianjin for Medical Epigenetics, Tianjin Key Laboratory of Medical Epigenetics, Department of Pharmacology, Tianjin Medical University, Tianjin 300052, P.R. China.
[54] School of Medical Imaging and Tianjin Key Laboratory of Functional Imaging, Tianjin Medical University, Tianjin 300052, P.R. China
[55] Department of Psychology, Institute of Psychiatry, Psychology and Neuroscience, King's College London, London SE5 8AF, United Kingdom
[56] Google, Inc., 1600 Amphitheatre Parkway, Mountain View, CA 94043, USA
[57] CAS Center for Excellence in Brain Science and Intelligence Technology, Chinese Academy of Sciences, Shanghai, 200031, P.R. China
[58] Population Neuroscience and Precision Medicine (PONS) LIN-Charite Research Group Dept. of Psychiatry and Psychotherapy, Charite, CCM, Humboldt University, Berlin, Germany and Institute for Science and Technology of Brain-inspired Intelligence (ISTBI), Fudan University, Shanghai, P.R. China.

[*] **Correspondence to:**

Professor Gunter Schumann, M.D., Ph.D., Centre for Population Neuroscience and Precision Medicine (PONS), Institute of Psychiatry, Psychology and Neurosciences, SGDP-Centre, 16 De Crespigny Park, London SE58AF, UK.

and
Professor Chunshui Yu, Ph.D., Department of Radiology, Tianjin Medical University General Hospital, No. 154, Anshan Road, Heping District, Tianjin 300052, China.


**Content:**

1. Supplementary Methods
2. Supplementary Results
3. Supplementary Figures S1-S5
4. Supplementary Tables S1-S6

### Supplementary Methods

#### CHIMGEN and IMAGEN project

Chinese Imaging Genetics (CHIMGEN) project led by Tianjin Medical University General Hospital initiated in 2015 and included 29 centers from 20 cities covering the most populous areas in China mainland. Until May of 2019, we have recruited 7,200 participants. The project was approved by the ethic committee of each center, written informed consent is obtained from each participant. Blood sample, environmental, structural and functional neuroimaging and cognitive assessment were prospectively collected from healthy Chinese Han participants of 18-30 ages to investigate genetic and environmental effects on brain and cognition.

Briefly, IMAGEN is the first European multisite and prospective project which aims to integrate different levels of environmental and biological mechanisms to identify biomarkers for developmental psychiatric disorders[1]. Comprehensive environmental factor, genetics, transcriptome, epigenetics, structural and functional neuroimaging, neurocognitive measure and mental health outcome are collected from more than 2000 14-year-old adolescents in 2009. Brain imaging measures were longitudinally assessed at age 14 years (Baseline (BL)) and 19 years (Follow-up 2 (FU2). Most of neurocognitive and mental health outcome longitudinally assessed at BL, FU1 (16 years) and FU2.

#### Geography data acquisition

In the CHIMGEN project, we recorded the precise residential addresses of each participant in each year from their birth to recruitment and the category of each place (1=rural, 2=town, 3=city) according to the National Bureau of Statistics (http://www.stats.gov.cn/tjsj/ndsj/renkoupucha/2000pucha/html/append7.htm). If some subjects moved several times in a year, the addresses where they live more than 6 months were recorded. Finally, 3336 participants with their complete living trajectory from the first stage of CHIMGEN project were included in the further analysis (Figure 1). In the IMAGEN project, the precise residential addresses/postcodes of 1083 participants at FU2 stage were recorded. Finally, 1025 participants were included in the further analysis because 58 participants move into other countries during IMAGEN BL and FU2 recruitment (from 2009 to 2015 year) (Figure 1). Finally, these addresses have been anonymized to 1km scaled coordinate (longitude and latitude) based on Google earth engine coordinate reference system for protecting personal information.

##### Remote sensing satellite data

Google earth engine (GEE) is an open access platform that makes hundreds of earth-observational satellite imagery and geospatial datasets with planetary-scale analysis available for researchers (https://earthengine.google.com/). Global Human Settlement Layer (GHSL)[2], Nighttime lights (NL)[3], Normalized Difference Vegetation Index (NDVI)[4], Normalized Difference Built-up Index (NDBI)[5], Normalized Difference Water Index (NDWI) [6] and global land cover mapping[7] were extracted from GEE to measure different urbanicity characteristics based on the acquired individual geography data from CHIMGEN and IMAGEN sample (Supplementary Table S3-S5).

##### Global Human Settlement Layer (GHSL)

Urbanicity was generally measured by dynamics of demographics in sociology[8,9]. GHSL is a framework to produce global spatial information about population and the physical size of settlements on the planet[2], of which population grid (GHSL-POP) is referred as the degree of urbanicity based on a classical theory of measuring urbanicity[8]. Product GHSL, Population Grid (P2016) for the epochs 1990-2000-2015 from GEE was applied in the participants from CHIMGEN and IMAGEN at 250m×250m resolution. Detailed information about the GHSL data can be found in https://ghsl.jrc.ec.europa.eu/index.php. As expected, population grid from GHSL in the town and city were much higher than those in rural for the epochs 1990-2000-2015 in CHIMGEN and IMAGEN (Figure S1).

**Nighttime Lights (NL)**

NL was used to detect visible and near-infrared emission sources at night, which could be applied to measure the prosperity or urbanicity of the neighborhood surroundings[3,10-12]. The band stable lights from Defense Meteorological Program (DMSP) Operational Line-Scan System (OLS) Nighttime Lights Time Series Version 4 product were extracted at 1km ×1km resolution from 1992.01.01 to 2014.01.01 based on the lifespan of the CHIMGEN and IMAGEN participants (https://developers.google.com/earth-engine/datasets/catalog/NOAA_DMSP-OLS_NIGHTTIME_LIGHTS).

**Normalized Difference Vegetation Index (NDVI)**

NDVI was used to assess outdoor surrounding greenness and generally applied to detect the urbanicity process[4]. Here, we used NDVI derived from product NOAA Climate Data Record (CDR) of Advanced Very High-Resolution Radiometer (AVHRR) NDVI at 5km×5km resolution from 1981.06.24 to 2017.10.05 based on the lifespan of the CHIMGEN and IMAGEN subjects (https://developers.google.com/earth-engine/datasets/catalog/NOAA_CDR_AVHRR_NDVI_V4).

**Normalized Difference Built-up Index (NDBI) and Normalized Difference Water Index (NDWI)**

As the name imply, NDBI and NDWI were used to assess built-up and water content of neighborhood surroundings, which were applied to map urbanicity[5,13]. Based on Band 1-7 from USGS Landsat 7 Collection 1 Tier 1 Raw Scenes (https://developers.google.com/earth-engine/datasets/catalog/LANDSAT_LE07_C01_T1) (Supplementary Table S4), NDBI was calculated as (B5-B4)/(B5+B4)[5] and NDWI was calculated as (B2-B4)/(B2+B4)[6] from 1999.01.01 to 2017.10.08, which both ranged from -1 to 1.

#### Global Land Cover Mapping

Global land cover mapping has been the most important variable on environmental changes and neighborhood surrounding resources[7,14,15]. Here Climate Change Initiative Land Cover dataset (CCI-LC) from European Space Agency platform was used to extract land cover classes from 1992 to 2015 (http://maps.elie.ucl.ac.be/CCI/viewer/). Briefly, CCL-LC aimed to make the best use of available satellite sensor data in order to provide an accurate percentage of land cover classes using supervised (machine learning) classification algorithm. The percentage here was defined as the number of pixels classified into a specific class divided by the total number of pixels within 1km radius centered at the anonymized home locations. There are 17 classes in CCL-LC including cropland, short tree, grassland, shrubland, snow and ice, water, urban and built-up, bareland and so on. In the present study, only land cover classes with mean percentage across all subjects above 1% were included in the further analysis. Finally, short trees (13.16%), cropland (64.21%), urban and built-up (31.31%) and water (7.92%) were selected in the CHIMGEN and short trees (10.23%), cropland (20.00%), urban and built-up (59.30%) and water (2.75%) were selected in the IMAGEN (Supplementary Table S3-S5).

#### Neuroimage acquisition

##### CHIMGEN project

In this study, only 1104 participants from the CHIMGEN Tianjin centers were included in the neuroimaging analysis. MRI data were acquired from 3.0 Tesla Discovery™ MR750 (General Electric®) and TrioTim Verio Skyra (Siemens®) scanners.

##### T1 weighted images acquisition

For the Discovery™ MR750 scanner, sagittal high-resolution three-dimensional T1-weighted imaging were acquired by Brain Volume imaging (3D-BRAVO) series with the following parameters: repetition time (TR)/echo time (TE) = 8.14/3.17 ms; inversion time (TI) = 450 ms; field of view (FOV) = 256mm × 256 mm; matrix = 256 × 256; flip angle (FA) = 12°, slice thickness (ST) = 1 mm; no gap; 188 sagittal slices. For the TrioTim Verio Skyra scanner, sagittal high-resolution three-dimensional T1-weighted imaging data were acquired by magnetization prepared gradient-echo (MP-RAGE) series with the following parameters: TR/TE = 2000/3.44 ms; TI = 900 ms; FOV = 256mm × 256 mm; matrix = 256 × 256; FA = 9°; ST = 1 mm; no gap; 192 sagittal slices.

**Diffusion tensor images (DTI) acquisition**

For the Discovery™ MR750 scanner, diffusion tensor images were acquired using spin-echo single-shot diffusion tensor echo plannar imaging (SE-SS-DT-EPI) sequence with the following parameters: TR/TE = 6000/61 ms; FOV = 256mm × 256 mm; matrix = 128 × 128; FA = 90°, ST = 3 mm; no gap; 50 axial slices; 64 diffusion gradient directions with b-value 1000 s/mm$^2$; 5 non-diffusion-weighted images (b = 0 s/mm$^2$); For the TrioTim Verio Skyra scanner diffusion tensor images were also acquired using the SE-SS-DT-EPI with the following parameters: TR/TE = 6500/89 ms; FOV = 256mm × 256 mm; matrix = 128 × 128; FA = 90°, ST = 3 mm; no gap; 50 axial slices; 64 diffusion gradient directions with b-value 1000 s/mm$^2$; 1 non-diffusion-weighted images (b = 0 s/mm$^2$).

**Resting-state fMRI acquisition**

For the Discovery™ MR750 scanner, resting-state functional MRI data were collected using gradient-echo single-shot echo-planar imaging (GRE-SS-EPI) with the following parameters: TR/TE = 2000/30 ms; FOV = 220mm × 220 mm; matrix = 64 × 64; FA = 90°; ST = 3 mm; gap=1mm; 40 interleaved transverse slices; 180 volumes. For the TrioTim Verio Skyra scanner, resting-state functional MRI data were also collected using GRE-SS-EPI with the following parameters: TR/TE = 2000/30 ms; FOV = 220mm × 220 mm; matrix = 64 × 64; FA = 90°; ST = 3 mm; gaP=0.99 mm; 36 interleaved transverse slices; 180 volumes. During the functional MRI scans, all subjects were instructed to keep still with their eyes closed, to think of nothing in particular, to stay as motionless as possible, and to not fall asleep.

**IMAGEN project**

In this project, 1724 and 1423 subjects from IMAGEN BL and FU2 were included in the neuroimaging analysis. Data were collected on eight 3 Tesla MRI scanners from different manufacturers (Siemens®, Philips®, General Electric®, Bruker®). In order to pool the data across sites, a phantom was scanned at each site to homogenize geometric distortions and signal uniformity. Moreover, several healthy volunteers were scanned at all sites to assess heterogeneity which was not captured by the phantom[1].

**T1 weighted images acquisition**

High resolution sagittal three-dimensional T1 weighted MP-RAGE images were acquired according to the ADNI protocol (http://adni.loni.usc.edu/methods/mri-analysis/mri-acquisition/) with the following standardized parameters across sites: TR/TE = 2300/2.8 ms; FA = 8°; matrix = 256×256; 160 volumes.

**DTI acquisition**

DTI were acquired using spin-echo single-shot diffusion tensor echo plannar imaging (SE-SS-DT-EPI) series with the following parameters: TR/TE = 6000/104 ms; FOV = 256mm × 256 mm; matrix = 128 × 128; FA = 90°, ST = 3 mm; no gap; 60 axial slices; 32 diffusion gradient directions with b-value 1300 s/mm$^2$.

**Resting-state fMRI acquisition**

Resting state fMRI scanning of the IMAGEN subjects were carried out at multiple sites with the following parameters: TR/TE = 2220/30ms; FOV = 218mm × 218mm; matrix = 64× 64; FA = 75°; ST = 2.4 mm; gap=3.4 mm; 164 volumes.

#### Neuroimage preprocessing

##### Gray matter volume (GMV) calculation

To exclude heterogeneity, the same preprocessing process of structural neuroimaging between CHIMGEN and IMAGEN cohorts were applied as following steps. A total of 1102 CHIMGEN subjects, 1724 IMAGEN subjects at BL and 1407 IMAGEN subjects at FU2 were finally included in the voxel-based morphometry (VBM) analysis. The GMV maps were calculated using the VBM8 implemented in Statistical Parametric Mapping software package (SPM8, http://www.fil.ion.ucl.ac.uk/spm). In the segmentation of VBM8, an adaptive Maximum A Posterior technique[16] and a Partial Volume Estimation[17] were used to estimate the fraction of each pure tissue type present in every voxel. After the structural images were segmented into gray matter (GM), white matter and cerebrospinal fluid, the individual's GM concentration map was normalized into the Dartel template in Montreal Neurological Institute (MNI) space (http://www.mni.mcgill.ca/). This template was derived from 550 healthy control subjects of the IXI-database (http://www.brain-development.org). In the modulated normalized process, we multiplied the individual's GM concentration map only by the non-linear determinants derived from the spatial normalization procedure. This step resulted in normalized GM density or relative GMV map for each subject. Here, the GMV of each voxel represented the fraction of GM present in each voxel, which preserved the local GM density while removing the confounding effect of variance in individual brain sizes. After that, we resliced the normalized GMV to a 1.5-mm cubic voxel. Finally, the GMV images were smoothed with a kernel of 8 × 8 × 8 $mm^3$ full width at half maximum. Then, the spatial pre-processing, normalized, modulated, and smoothed GMV maps were used for further analysis.

Finally, 831 CHIMGEN subjects, 810 IMAGEN BL subjects and 791 IMAGEN FU2 subjects with available quantified both GMV data and remote sensing satellite features were included in the further analysis (Table 1).

**Tract based spatial statistics (TBSS) calculation**

A total of 1515 CHIMGEN subjects, 1958 IMAGEN subjects at BL and 1322 IMAGEN subjects at FU2 were finally included in the tract based spatial statistics (TBSS) analysis[18]. DTI data from CHIMGEN and IMAGEN project were both pre-processed using software from the FMRIB's Software Library (FSL v6.0.1) toolbox (www.fmrib.ox.ac.uk/fsl)[19] in the following manner: An affine registration was applied to the first b0 image for head motion and eddy current correction. Brain extraction was carried out using BET. Diffusion tensor fitting was then used to obtain fractional anisotropy (FA) maps for each subject. All participants' FA data was aligned into a common space using the non-linear registration tool FNIRT, using a b-spline representation of the registration warp field. The mean was then taken across all FA maps to create an FA averaged image. This map was then 'thinned' to create a mean FA skeleton, which was then thresholded at FA>0.2, keeping only the major white matter tracts. Each subject's aligned FA data was then projected onto the mean skeleton. A further 10 scans were not used due to masking or normalization issues in TBSS. We then used these skeletonized maps in all subsequent analyses.

Finally, 849 CHIMGEN subjects, 940 IMAGEN BL subjects and 724 IMAGEN FU2 subjects with available quantified both TBSS data and remote sensing satellite features were included in the further analysis.

**Resting-state fMRI preprocessing**

**CHIMGEN project**

The fMRI data of the 1104 CHIMGEN subjects were preprocessed using the Data Processing

Assistant for Resting-State fMRI (DPARSFA) in the following steps[20].

(1) The first 10 volumes from each subject were discarded to allow the signal to reach equilibrium and to allow the participant to adapt to the scanning noise. The remaining 170 volumes were corrected for the acquisition time delay between slices;

(2) Rigid realignment was then performed to estimate and correct the motion displacement. Six subjects were excluded from further analysis because their fMRI data had a maximum displacement in one or more of the orthogonal directions (x, y, z) of > 2 mm or a maximum rotation (x, y, z) > 2.0°. Thus, a total of 1098 subjects' fMRI data was included in the further analysis;

(3) All data were then spatially normalized to the standard echo planar image (EPI) template and resampled to a voxel size of 3 × 3 × 3 $mm^3$;

(4) The normalized data were smoothed by a Gaussian kernel of 4 × 4 × 4 $mm^3$;

(5) After normalization, several nuisance covariates (six estimated motion parameters and average BOLD signals of the whole brain and ventricular and white matter regions) were removed from the data using a regression analysis.

Finally, 827 CHIMGEN subjects with available quantified both fMRI data and remote sensing satellite features were included in the further analysis

##### IMAGEN project

The fMRI data of the 381 subjects from IMAGEN BL and 1066 subjects from IMAGEN FU2 were collected. Of these scans, 158 were not used, either because over 5% of scans in that subject exhibited artifacts of some kind, or if over 5% of volumes showed a fractional displacement of over 0.5mm. Preprocessing of resting-state data was performed with routines from FSL v5.0.9 and Advanced Normalization Tools (ANTs v1.9.2).

(1) Motion correction was carried out, applying a rigid body registration of each volume to the middle volume (FSL MCFLIRT);

(2) Non-brain tissue was removed (FSL BET);

(3) Spatial smoothing was applied using a Gaussian kernel of $4 \times 4 \times 4$ $mm^3$;

(4) Independent component analysis (FSL MELODIC) was run for each data set. Artifact components were identified using an automatic classification algorithm, and subsequently regressed from the data (ICA-AROMA v0.3)37). ICA-AROMA has been shown to be superior to other motion correction procedures, for the purposes of removing motion related artifacts;

(5) The resulting cleaned data set was de-trended (up to a third degree polynomial);

(6) Co-registration to a high-resolution T1 image (FSL FLIRT using the BBR algorithm), and normalization to 2mm isotropic MNI standard space (ANTs) was carried out;

(7) To further clean the data of physiological noise using CompCorr procedure, we created white matter (WM) and cerebrospinal fluid (CSF) masks by taking the mean of the WM and CSF segmentations from the VBM analysis, and thresholding them at 0.95, we then resliced these maps into the same space as the rsfMRI data. We then extracted timecourses from voxels within these regions and took the first three principal components of this signal for both WM and CSF maps. These six principal component signals should represent non-neuronal signal. We then regressed this non-neuronal signal from voxel timecourses across the rest of the brain.

(8) Lastly, preprocessed and normalized resting-state data sets were resliced to 3mm isotropic voxels.

Finally, 156 IMAGEN subjects at BL and 614 IMAGEN subjects at FU2 with available quantified both fMRI data and remote sensing satellite features were included in the further analysis

#### Identification of resting state network

In the CHIMGEN and IMAGEN sample, group independent component analysis (GICA) was used to decompose all the resting-state fMRI data into independent components (ICs) to construct brain functional network using the GIFT software (http://mialab.mrn.org/software/gift/index.html, version 4.0b), which included data reduction, ICA and back reconstruction[21]. Data reduction was used to reduce the size of the subject's fMRI data using the principal components analysis. Two data reduction steps were adopted. After each subject's fMRI data was reduced, the subjects were concatenated into one group and put through another data reduction step. ICA algorithm was then applied to the reduced dataset to identify ICs. The number (n = 40) of ICs was automatically estimated using the minimum description length (MDL) criterion[22]. ICASSO toolbox was used to determine the reliability of ICA algorithm. Specifically, ICA was run 100 times to obtain the final integrated output. The subject-specific time courses and spatial maps were back-reconstructed by a dual-regression method. A linear spatial regression was applied to the group-level spatial maps and individuals' fMRI datasets to calculate matrix describing time courses for each component of each subject[21]. A linear temporal regression was then applied to these time-course matrices and individuals' fMRI datasets to estimate subject-specific spatial maps [23]. For each subject-specific spatial component, the value of a voxel represents the relation of the time courses between this voxel and the subject-specific component. To improve the normality of the data, we scaled the spatial component maps to z-scores[21,23]. Using GICA of estimated 40 network components[21,22], we identified 17 resting-state networks (RSN) related to various cognitive and sensory-motor processes[24] that were similar in the CHIMGEN and IMAGEN samples (Figure 3A).

**Neuropsychological assessment**

To test the overall exposure effect of urbanicity on behaviors, we included different dimension of neuropsychological assessment and mental health features. Only 621 subjects from CHIMGEN, 637 subjects from IMAGEN FU1 and 578 subjects from IMAGEN FU2 with available quantified assessment data and remote sensing satellite features were included in the further analysis.

**Verbal learning memory**

Here the California Verbal Learning Test (CVLT) was used to test episodic verbal learning and memory, which demonstrates sensitivity to a range of clinical conditions. Briefly, the experimenter would read a list of 16 nouns words loudly every second for five sessions in a fixed order. After each session, the subjects were asked to try their best to recall the words in any order. Then we will record the numbers of correct words in immediate free recall, short-term free recall, short-term clue recall, long-term free recall, long-term clue recall and long-term recognition sessions.

**Working memory**

Here classic letter N-back test (1-back and 3-back) were applied to measure working memory. In the letter 1-back task, the subjects were required to match the current stimulus by the previous one in the sequence of stimuli. In the letter 3-back task, the subjects were required to match the current stimulus by the one from 2 steps earlier in the sequence of stimuli. Both 1-back and 3-back task have only one block including 60 trails. Each stimulus was presented for 200 ms and the stimulation interval is 1800 ms. The participants were instructed to respond as fast and accurately as possible after the presentation of each stimulus. Before the experiment, the subjects were given one practice test run of the 1-back task. Only the subjects with correct rate more than 75% were allowed to formal task. If not, they have to do the test task again. E-Prime 2.0 software (Psychology Software Tools) was used to present the stimuli and collect the results. We recorded the numbers of correct rejection stimuli (cr), hits stimuli (h), misses stimuli, false alarms stimuli, no response stimuli and finally calculate the correct rate ((cr+h)/60).

**Information processing speed**

Here Symbols Digit Modality Test (SDMT) was applied to test attention and speed of processing ability. Using a reference key, the subjects were required to pair specific numbers with given nine geometric figures after testing the first 10 items as fast and accurate as they can. Finally, we record the numbers of correctly and incorrectly filled digits in the 90 seconds. The highest score was 110 points.

##### Social cognition

In the CHIMGEN, a ball tossing game with a 2×2 factorial design was applied to test perspective taking (first-person vs. third-person perspective) and agency (active vs. passive) of social cognition. The subjects were required to perform active and passive tasks from two different perspectives. From the third-person perspective (3PP), three virtual characters (red, green, and blue) appeared on the screen to perform a ball tossing game. They have to perform the two tasks from the blue characters' perspectives instead of from their own. In the active task, subjects were instructed to throw the ball to the red characters by pressing the corresponding key "F" or "J" (left or right) when they were in possession of the ball. In the passive task, one of the two characters were in possession of the ball, and subjects have to judge the ball's position in relation to themselves (left or right side) with a button press "F" or "J"; From the first-person perspective (1PP), the visional field of the subject was consistent with the forward vision of the blue character. Only one hand of the blue man was displayed on the screen without the body. They were also asked to perform the active and passive tasks. The whole task includes 72 trials from four blocks in the order of 1PP, 3PP, 3PP and 1PP block. Each block contains 18 trails including 6 active tasks and 12 passive tasks. The location of the subject, the player in possession of the ball and the relative position of the red man are all pseudo-random. The schematic representation of the ball tossing task design was shown in Figure S2.

The participants were instructed to respond as fast and accurately as possible after the presentation of each trail. Before the experiment, the participants were given 2 practice runs of the task. E-Prime 2.0 software was also used to present the stimuli and collect the results. We recorded active reaction time in the 1PP and 3PP (1PP_ACT_RT and 3PP_ACT_RT), active accuracy in the 1PP and 3PP (1PP ACT_ACC and 3PP_ACT_ACC), passive reaction time in the 1PP and 3PP (1PP_PAS_RT and 3PP_PAS_RT), passive accuracy in the 1PP and 3PP (1PP PAS_ACC and 3PP_PAS_ACC). The perspective taking and agency performance were evaluated by 4 measures: reaction time in perspective taking (PT_RT) was defined as sum of 3PP_RT minus 1PP_RT (PT_RT = (3PP ACT_RT + 3PP PAS_RT) - (1PP ACT_RT+1PP PAS_RT) ); accuracy in perspective taking (PT_ACC) was defined as sum of 3PP_ACC minus 1PP_ACC (PT_ACC = (3PP ACT_ACC + 3PP PAS_ACC) - (1PP ACT_ACC+1PP PAS_ACC)); reaction time in agency (AG_RT) was defined as sum of PAS_RT minus ACT_RT (AG_RT = (3PP PAS_RT + 1PP PAS_RT) - (3PP ACT_RT+1PP ACT_RT); accuracy in agency (AG_ACC) was defined as sum of PAS_ACC minus ACT_ACC (AG_ACC = (3PP PAS_ACC + 1PP PAS_ACC) - (3PP ACT_ACC+1PP ACT_ACC) in the further anlaysis. Ten subjects were excluded due to lack of the ball tossing game task results.

In the IMAGEN FU1 and FU2 sample, Interpersonal Reactivity Index (IRI) questionnaire was used to test dispositional empathy comprising four separate but related conducts: perspective taking, fantasy, empathic concern and personal distress.

**Executive control**

Here Go/No-Go tests were used to measure participants capacity for sustained attention and response control. The subjects were required to perform an action when the current stimulus was different from the previous one (e.g., press a button - Go trail) and inhibit that action when they were match (e.g., not press the same button - No-Go trail) in the sequence of stimuli. The whole task included two blocks with 21 Go trails and 189 No-Go trails in each block. The participants were instructed to respond as fast and accurately as possible after the presentation of each trail. Before the experiment, the participants were given 1 practice run of the task. E-Prime 2.0 software was used to present the stimuli and collect the results. We recorded accuracy in Go trial and No-Go trails, respectively.

##### Mental health

In the CHIMGEN, Beck Depressive Inventory (BDI) was used to measure severity of depression. And State-Trait Anxiety Inventory (STAI）was used to measure state anxiety (anxiety about an event) and trait anxiety (anxiety level as a personal characteristic). In the IMAGEN, Ruminating Scale (RSQ) was used to measure the frequency of cognitions and behaviors of subjects during periods of depressed mood, which was only available at FU2 sample. And Anxiety Screening for Composite International Diagnostic Interview (CIDI-DIA) and The Development and Well-Being Assessment Interview (DAWBA) were applied to measure their anxiety state, which was available in FU2 sample.

#### Statistical analysis

##### Remote sensing satellite data

To test the latent urbanicity behind the satellite features, confirmatory factor analysis (CFA) was applied to the satellite features using R package *lavaan* (https://cran.r-project.org/web/packages/lavaan)[25]. With CFA, one can estimate how well the measured satellite variables represent the latent urbanicity. Firstly, eight satellite features related to urban characteristics including NL, NDVI, NDWI, NDBI, percentage of cropland, percentage of urban build up, percentage of short tree and percentage of water were correlated with each other to identify the promising measured variables. Only the closely correlated features ($r > 0.4$) were included in the CFA model to construct a factor for urbanicity in China and Europe respectively, which we termed 'UrbanSat' (Supplementary Table S6). To consolidate our foundation of using satellite features to measure urbanicity, the UrbanSat measure was correlated with ground-level population grid data from GHSL-POP in China and Europe respectively, which was typically used to map urbanicity process in sociology[8,9].

##### Neuroimaging data

##### VBM and TBSS analysis

The voxel-wise multiple regression analysis was performed to identify brain regions whose GMV or FA were significantly correlated with UrbanSat using Statistical Parametric Mapping software package (SPM12, http://www.fil.ion.ucl.ac.uk/spm), while controlling for the age, gender and neuroimaging sites. For TBSS analysis of the FA, the threshold-free cluster enhancement (TFNCE) option in the permutation-testing tool (permutations = 5000) in the FSL software was used to test statistical significance[26]. Multiple comparisons were corrected using a voxel-level family-wise error (FWE) method ($P$<0.05, cluster size >100 voxels). The GMV or FA of brain regions with significant correlations with the UrbanSat measure were extracted for further analysis.

##### Within-network FNC

The 17 ICA components were entered a random-effect one-sample t-test to generate sample-specific spatial maps (FWE correction, $T$=8). Then a voxel-wise multiple regression was applied to test the correlation between UrbanSat and FNC within the mask of 17 brain networks, respectively. Multiple comparisons were corrected using a voxel-level FWE method ($P$<0.05, cluster size >100 voxels). The FNC of brain region within the brain networks showed significant correlation with UrbanSat were extracted for the further analysis.

##### Between-network FNC

*Pearson* correlation was calculated for the ICA timecourses between each pair of 17 networks for each subject and then converted $r$ value to $z$ values to improve normality. Each $z$ value represents the between-network FNC. Finally, partial correlation was applied between UrbanSat and FNC of 136 (17×16/2) between-networks, controlling for the effect of age, gender and neuroimaging sites. The permutation test with 10,000 times ($P$ < 0.05) was used to correct multiple comparisons.

**Susceptible developmental windows for the correlate of UrbanSat with brain**

If the above results were significant, independent voxel-wise multiple regression analysis was performed to test susceptibility windows of the correlation of UrbanSat with brain for each age sliding window group of CHIMGEN sample (FWE correction, voxel $P<0.05$, cluster size > 100 voxels). For example, the second age sliding window group refers to the subjects with UrbanSat of 1, 2 and 3 years old. To directly compare the statistic differences between the correlate of UrbanSat with brain feature in each age sliding window, R package *cocor 1.0-1* was used for comparing non-overlapping correlations procedure[27]. Finally, the significant correlate of UrbanSat with brain in different age sliding windows would be replicated in the IMAGEN sample.

**Neuropsychological test and mental health data**

Partial correlation analysis was applied to test the correlation between UrbanSat and each neuropsychological domain and mental health while controlled for age, gender and sites in CHIMGEN. Bonferroni correction ($P < 0.05$) was applied in multiple comparisons. Finally, the significant correlate of UrbanSat with behavior would be replicated in the IMAGEN sample.

**Mediation analysis**

The mediation analysis was performed to test whether the brain feature of each significant region mediated the correlation between UrbanSat and behaviours[28,29]. The UrbanSat was defined as an independent variable, the brain features of each significant region as a mediator variable, and the significant behavior as a dependent variable. The first step is to confirm that the independent variable (UrbanSat) is a predictor of the dependent variable (cognitive and mental health outcome), which is known as the direct effect. The second step is to confirm that the independent variable (UrbanSat) is a predictor of the mediator (brain). The third step is to confirm that the mediator (brain) is a predictor of the dependent variable (cognitive and mental health outcome), while controlling for the independent variable (UrbanSat). The indirect effect is the product of path coefficients of the last two steps. Then the bootstrapping method is used to assess the significance of the mediation effect. After 5000 bias-corrected bootstrapping, we could estimate the distribution of the indirect effect and calculate its 95% confidence intervals (CI). If zero does not fall between the resulting 95% confidence interval of the bootstrapping method, we could confirm the existence of a significant mediation effect ($P < 0.05$). Finally, the significant mediation pathway was replicated in the IMAGEN datasets.

#### Supplementary Results

##### The relation of population density with brain GMV

A voxel-wise multiple regression analysis showed that average Chinese population grid from 1990 to 2015 were positively correlated with cerebellar vermis volume (MNI coordinate: x=0, y=-57, z=-12; 273 voxels; F value peak = 5.78； Figure S3A) and negatively correlated with bilateral medial superior frontal cortex volume in the whole brain (MNI coordinate: x=0, y=34.5, z=39; 1377 voxels; F value peak = -6.19； Figure S3B) while controlled age, gender and sites (FWE correction, voxel wise $P$<0.05, cluster size>100 voxels). We observed similar results in the effect of UrbanSate on brain GMV, thus validating the effect of urbanicity obtained by satellite measures on brain volume.

##### Vulnerable periods analysis between longitudinal UrbanSat and brain GMV

We investigated potential susceptibility periods during which exposure to urbanicity may have the strongest influence on brain GMV. In the CHIMGEN participants, for whom geoposition data were available since birth, we created age sliding windows by averaging UrbanSat for each subject over a period of three years and measured correlations between the exposure to urbanicity in each sliding window and GMV in young adulthood (23.81±0.82 years). While we found no significant effect of UrbanSat on brain GMV from age 1 to 3 years, from age 4 years, UrbanSat was negatively correlated with the left mPFC volume and positively correlated with cerebellar vermis volume. (FWE-correction, voxel wise $P$<0.05, cluster size>100 voxels). This correlation lasts until age 16 years, without significant effect of UrbanSat on either left mPFC or cerebellar vermis volume from age 17 to 25 years. From age 11 to 14 years, the negative correlation between UrbanSat and the left mPFC volume was significantly greater than in other age windows. The difference to the adjacent age bands was significant at $P$=0.012 compared to age 10 years and $P=2.36\times10^{-5}$ compared to age 15 years, suggesting a susceptibility window for the effect of urbanicity on left mPFC volume in adolescence (Figure 2B). The positive correlation between UrbanSat and the cerebellar vermis volume was highest from age 4 to 7 years. The difference to the adjacent age bands was significant at $P$=0.016 compared to age 8 years, indicating a susceptibility window for the effect of urbanicity on the cerebellar vermis volume in childhood (Figure 2B).

**The relation of UrbanSat with within-networks FNC**

Resting-state FNC can reflect distinct cognitive or emotional processes that are influenced by individual developmental trajectories[30] and environmental factors[31], potentially resulting in dysfunctional connectivity associated with mental disorders[32]. Using Group-Independent-Component-Analysis (GICA) to estimate 40 network components[21,22], we identified 17 resting-state networks in both samples that were related to various cognitive and sensory-motor processes[24] (Figure 3A). To investigate the shared and distinct relations of urbanicity with brain activity of cognitive processes in China and Europe, we tested the association of UrbanSat with FNC of 17 within-network and 136 (17*16/2) between-networks.

In the CHIMGEN (N=827), a voxel-wise multiple regression analysis controlling for age, gender and sites revealed a negative correlation with FNC within the left mPFC of the aDMN (MNI-coordinate: x=6, y=33, z=57; 130 voxels; $F$-value peak=-4.96; Figure 3B) and a positive correlation of the UrbanSat with FNC within the left lingual gyrus of the medial visual network (mVN) (MNI-coordinate: x=-6, y=-84, z=-6; 144 voxels; $F$-value peak=5.21; Figure 3B) (FWE-correction, voxel $P$<0.05, cluster size>100 voxels). In the voxel-wise multiple regression analysis of IMAGEN at 19 years (N=614) controlling for age, gender and sites, UrbanSat showed a negative correlation with FNC within the left mPFC of aDMN (MNI-coordinate: x=-3, y=63, z=6; 106 voxels; $F$-value peak=4.83; Figure 3D), but no significant correlation with FNC within the left lingual gyrus of the mVN (FWE-correction, voxel $P$<0.05, cluster size>100 voxels). The findings in CHIMGEN and IMAGEN were consistent with the gray matter volume changes, indicating that the observed correlations between UrbanSat and the left mPFC were present in both structural and functional modalities

To identify a potential susceptibility period for urbanicity-related within-network connectivity of the aDMN and mVN, we carried out a voxel-wise multiple regression in each age sliding window controlled for age, gender and sites (FWE-correction, voxel wise $P<0.05$, cluster size>100 voxels). We found significant negative correlation of UrbanSat on FNC within the aDMN from age 3 years until age 16 years. From age 13 to 15 years, the negative correlation between UrbanSat and the FNC within the aDMN was significantly greater than at age 12 years ($P=0.010$) and at age 16 years ($P=6.16\times10^{-5}$) (Figure 3C). We confirmed in the IMAGEN participants at age 14 years the negative correlation with the FNC of the left mPFC within the aDMN (BA 10, MNI-coordinate: x=0, y=60, z=24; 104 voxels; *F* value peak=-5.61; Figure 3E) (FWE-correction, voxel wise $P<0.05$, cluster size>100 voxels). Together, our data suggest a susceptibility window for the correlation of urbanicity and FNC within the aDMN in adolescence. We also found a significant correlation of UrbanSat with FNC within the LG in the mVN only in childhood from age 4 to 11 years (Figure 3C). The positive correlation between UrbanSat and FNC within the mVN was highest from age 5 to 8 years. The difference to the adjacent age bands was significant compared to age 4 years ($P=0.026$) and to age 9 ($P=2.60\times10^{-4}$), indicating a susceptibility window for the effect of urbanicity on the FNC within the mVN in childhood (Figure 3C).

**The relation of UrbanSat with between-networks FNC**

The resulting 136 between-network FNCs of CHIMGEN were highly correlated with those of IMAGEN at age 19 years ($r=0.55$, $P=2.54\times10^{-12}$) and IMAGEN at age 14 years ($r=0.28$, $P=0.001$). As expected, the averaged 136 between-network FNCs of IMAGEN were highly correlated at age 14 years and 19 years ($r=0.60$, $P=1.15\times10^{-14}$) (Figure S4). These results demonstrated overall similarity of resting-state activity between CHIMGEN and IMAGEN. UrbanSat correlates with 42 between-networks FNCs in CHIMGEN (Figure 4A), and 19 between-networks FNCs in IMAGEN participants at age 19 (Figure 4B) ($P<0.05$ after 10,000 permutations), suggesting a greater sensitivity to urbanicity in the Chinese CHIMGEN sample compared to the European IMAGEN sample. We found 7 between-networks FNCs present in both samples, indicating a proportion of shared correlations between urbanicity and between-networks FNCs in Chinese and European samples. The shared between-networks FNCs involved the aDMN and several brain networks, including aDMN-ECN, aDMN-CBN, aDMN-rFPN (Table 2). In addition, we found shared between-network connectivity of rFPN -LN, rFPN-lFPN, AN-VN and AN-vSMN (Table 2). Adolescence was the period of greatest susceptibility for four between-networks FNCs: aDMN-ECN ($r$=0.165, $P$=3.25×$10^{-5}$), aDMN-CBN ($r$=0.130, $P$=1.09×$10^{-3}$), aDMN-rFPN ($r$=0.133, $P$=8.34×$10^{-4}$) and rFPN-lFPN ($r$=0.094, $P$=0.018) (Figure 4E and Table 2); Childhood showed highest correlations with 3 between-networks FNCs: AN-vSMN ($r$=0.131, $P$=1.04×$10^{-3}$), AN-VN ($r$=0.097, $P$=0.016) and rFPN-LN ($r$=0.130, $P$=1.09×$10^{-3}$) (Figure 4E and Table 2).

Whereas we found 19 between-networks FNCs associated with UrbanSat in IMAGEN FU2 at age 19, we detected 27 significant between-networks FNCs in the same participants at age 14 ($P$<0.05 after 10,000 permutations) (Figure 4C): Six between-networks FNCs were shared by IMAGEN participants at age 19 (pDMN-vAN, aDMN-CBN, aDMN-rFPN, aDMN-ECN, rFPN-LN, and rFPN-lFPN). Five between-networks FNCs were shared with IMAGEN and CHIMGEN (aDMN-CBN, aDMN-rFPN, aDMN-ECN, rFPN-LN, and rFPN-lFPN) (Figure 4D and Figure S5). These results indicate that between-networks FNCs relate to urbanicity in both shared and distinct ways during brain development and in different socio-cultural and geographic locations. Shared urbanicity-related between-networks FNCs include the aDMN and FPN, which may underlie self-referential thoughts[33] and executive control[34] during brain development.

#### The relation of UrbanSat with behavior

We investigated whether UrbanSat was correlated with measures of cognition and mental health-relevant behavior, in particular symptoms of depression and anxiety. Neuropsychological measures in CHIMGEN include verbal learning memory, working memory, information process speed, executive control and social cognition (perspective taking and agency). Of these domains, only social cognition, namely perspective taking, a measure of perceiving a situation or understanding a concept from an alternative point of view[35], was significantly positively associated with UrbanSat (accuracy: $r$=0.124, $P$=0.002; reaction time, $r$=-0.245, $P$= $6.10\times10^{-7}$; Figure S6 and Table 3) (Bonferroni corrected). The correlation between UrbanSat and perspective taking performance was strongest during adolescence (age 13 to 16 years in accuracy and 14 to 18 years in reaction time) (Figure 4F). The difference to the adjacent age bands was significant in accuracy ($P$=0.026 compared to age 12 year and $P$=$1.45\times10^{-4}$ compared to age 17 years) and in reaction time ($P$=$6.58\times10^{-5}$ compared to age 13 years and $P$=$1.16\times10^{-6}$ compared to age 19 years). We confirmed the positive correlations between UrbanSat and perspective taking in IMAGEN at age 16 years ($r$=0.103, $P$=0.009), the earliest age these data were available (Figure S6 and Table 4). In the IMAGEN at age 19 years, we observed a trend-level significant correlation between UrbanSat and perspective taking ($P$=0.056) (Figure S6 and Table 4).

In CHIMGEN, UrbanSat was correlated with number of symptoms in the Beck Depressive Inventory (BDI) ($r$ = 0.209, $P$=$1.44\times10^{-5}$) and state anxiety ($r$ = 0.132, $P$=0.001) (Bonferroni correction, $P$ < 0.05) (Figure S6 and Table 3). The correlations between UrbanSat with BDI and state anxiety were strongest in adolescence; in BDI at age 12 to 16 years ($P$=0.016 compared to age 11, and $P$=$3.62\times10^{-5}$ to age 17) (Figure 4F); in state anxiety at age 14-17 ($P$=0.036 compared to age 13 and $P$=0.011 to age 18) (Figure 4F). In IMAGEN, we validated the correlation between UrbanSat and depressive symptoms at age 19 ($r$=0.118, $P$=0.004) using the Ruminating Scale Questionnaire (RSQ). There was no association between UrbanSat and anxiety at any timepoint in IMAGEN (Figure S6 and Table 4).

#### Supplementary Figures

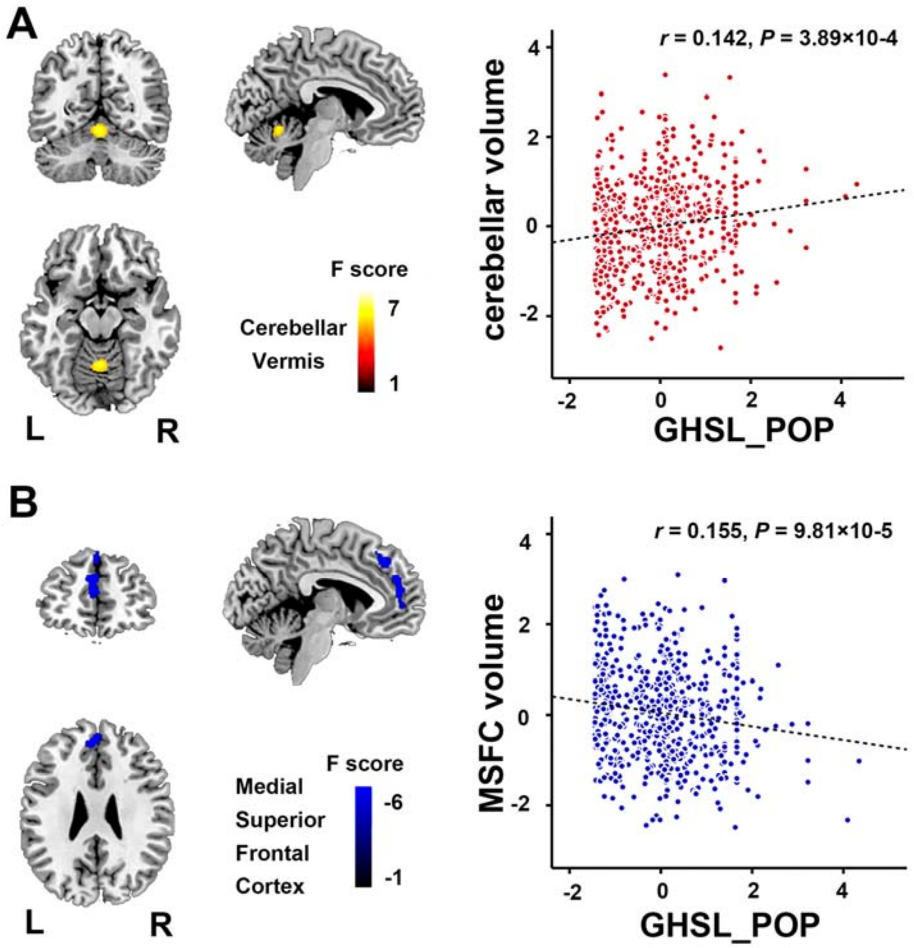


***Figure S1. Voxel wise correlation between GHSL_POP and GMV across the whole brain***. *A-B. In the CHIMGEN sample (N=831), there was a significant positive correlation of the average population grid (measured by GHSL_POP before age 18 years) with cerebellar vermis volume (A) and a significant negative correlation with left medial superior frontal cortex volume (B) (FWE correction, voxel P<0.05); GHSL, Population density grid from Global Human Settlement Layer; L, left; R, right.*

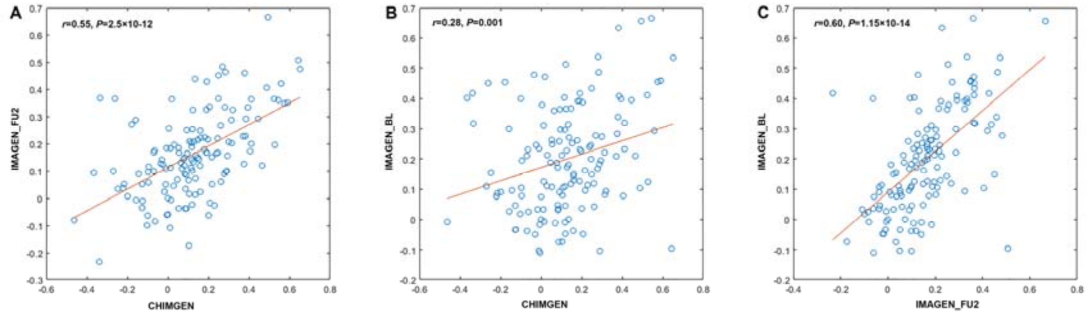


***Figure S2. Correlation maps between 136 averaged between-network FNC from CHIMGEN and IMAGEN FU2 (A), CHIMGEN and IMAGEN BL (B) and IMAGEN BL and FU2 (C).*** *These maps demonstrated overall similarity of resting-state activity between CHIMGEN and IMAGEN. FNC, functional network connectivity.*

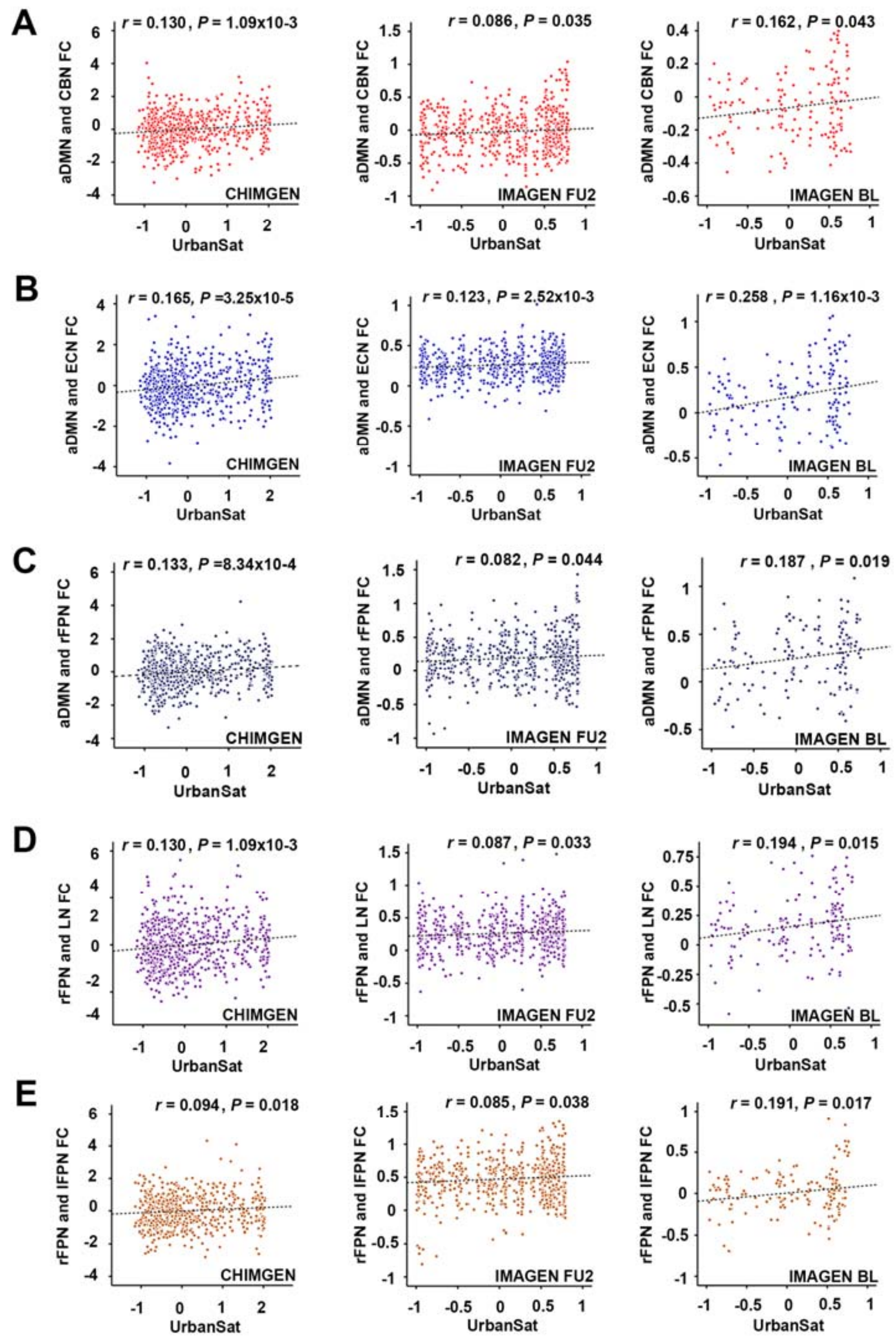


***Figure S3. The correlation of UrbanSat with five overlapped between-network FNC.*** *A-E. UrbanSat was significantly correlated with between-network aDMN-CBN (A), aDMN-ECN (B) and aDMN-rFPN (C), rFPN-LN (C), rFPN-lFPN (D), in the CHIMGEN, IMAGEN FU2 and BL sample after 10,000 permutations. aDMN, anterior defaut mode network; CBN, cerebellar network; ECN, executive control network; FNC, functional network connectivity; LN, language network; lFPN, left frontal parietal network; rFPN, right frontal parietal network.*

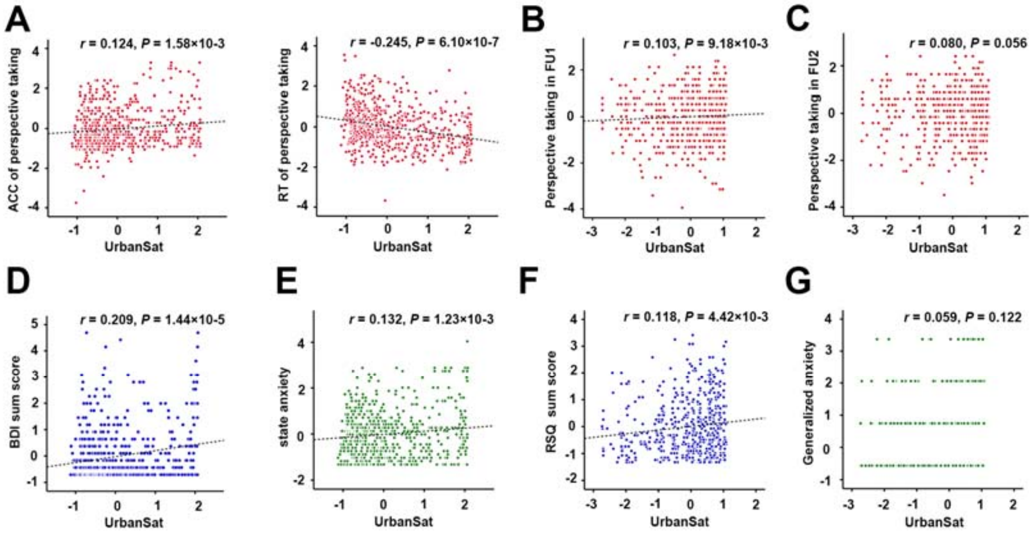


***Figure S4. The correlation of UrbanSat with perspective taking performance, depressive severity and state anxiety.*** *A. In the CHIMGEN sample, UrbanSat was significantly positively correlated with accuracy and negatively correlated with reaction time of perspective taking performance (Bonferroni correction, P<0.05); B-C. This result was replicated in the IMAGEN BL sample(B). In the IMAGEN FU2 sample, trend-level significance was observed (P=0.056) (C); D-E. In the CHIMGEN sample, UrbanSat was significantly positively correlated with BDI score (D) and state anxiety (E) (Bonferroni correction, P<0.05); F-G. The correlation of urbanicity on depressive severity (F), but not on generalized anxiety (G) was replicated in the IMAGEN FU2 sample at 19 years.*

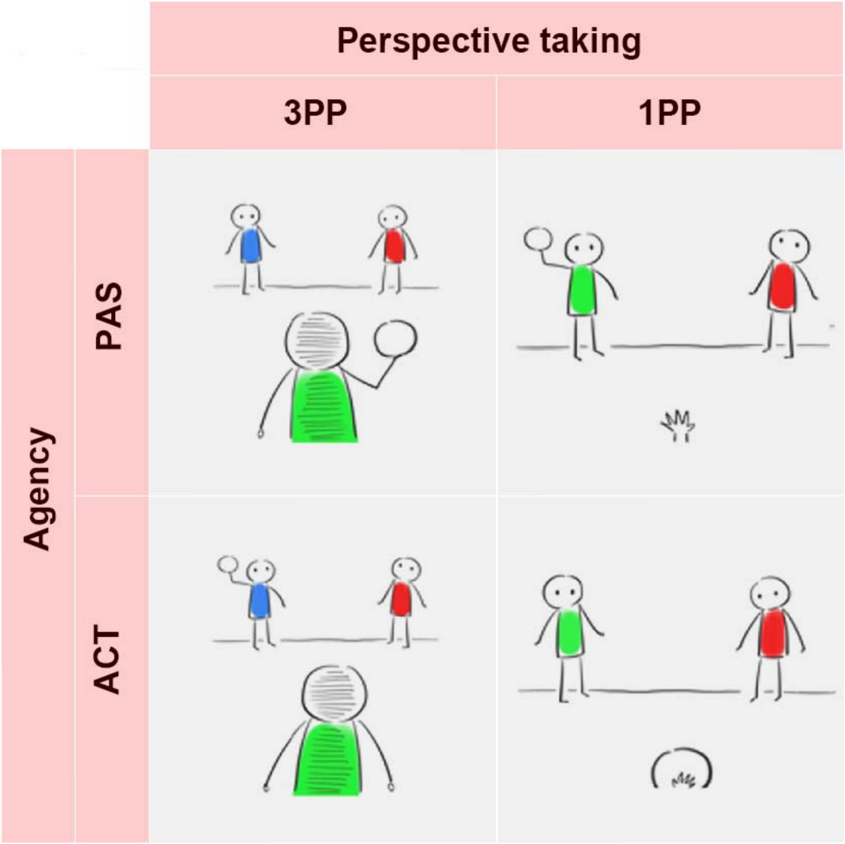


746
747 ***Figure S5. The schematic representation of the ball tossing task design, which measured perspective taking and agency performance.***
749
750
751
752

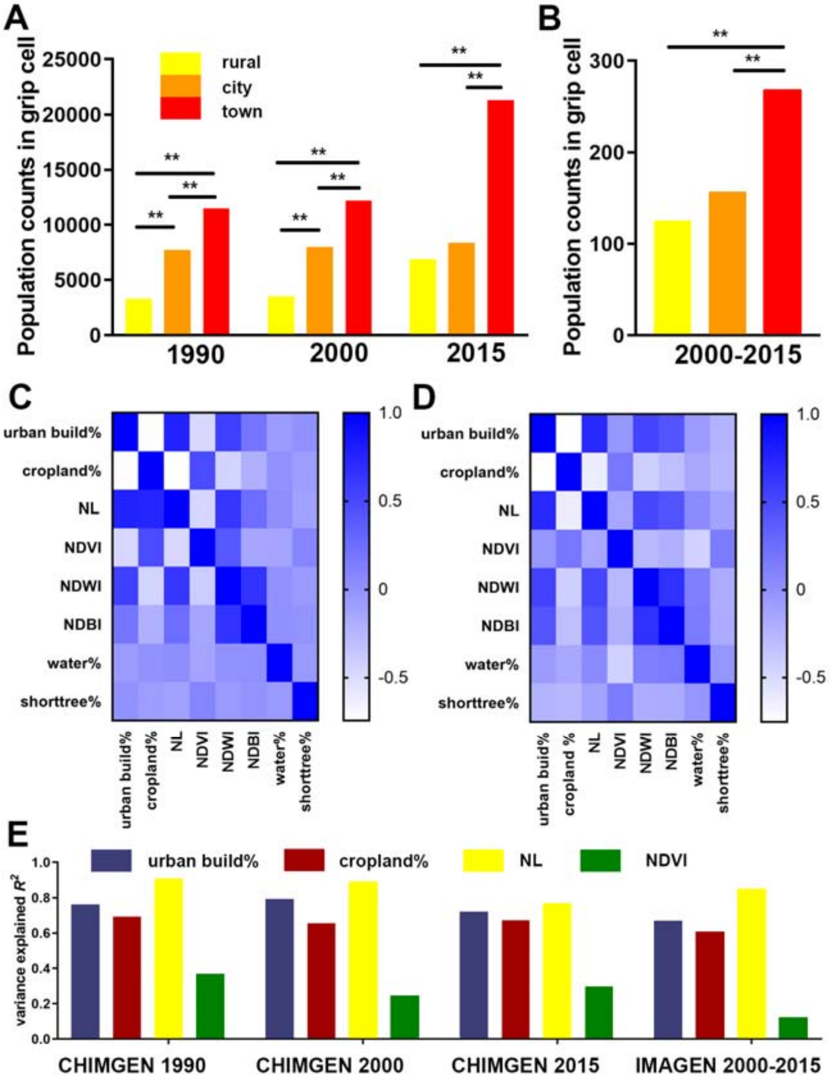


***Figure S6. Confirmatory factor analysis in CHIMGEN and IMAGEN.*** *A. Population grid distribution in rural, town and city for the years 1990, 2000 and 2015 in CHIMGEN sample; B. In IMAGEN sample, we selected only participants who remained at the same address (geoposition) between 2009 and 2015. We therefore used the average population grid between the years 2000 and 2015; C-D. Correlation matrix between eight remote sensing satellite features in CHIMGEN (N=3305) (C) and IMAGEN (N=1205) (D) sample. Only closely correlated features (r>0.4) (percentage of urban build-up, percentage of cropland, nightlights [NL] and NDVI) were included in the confirmatory factor model; E. Variance explained $R^2$ of each promising satellite feature for the Chinses and European UrbanSat.*

#### Supplementary Tables

**Table S1. Demographics of CHIMGEN and IMAGEN sample**

| Project | N | Gender (M/F) | Mean age (SD) |
|---|---|---|---|
| **CHIMGEN** | 831[a] | 391/440 | 23.81 (0.82) |
| **IMAGEN BL** | 810[a] | 362/448 | 14.05 (0.75) |
| **IMAGEN FU1** | 635[b] | 277/358 | 16.12 (0.76) |
| **IMAGEN FU2** | 791[a] | 369/422 | 19.00 (0.71) |

BL, baseline; FU1, follow up 1; FU2, follow up 2; SD, standard deviation

[a] Numbers of subjects with available UrbanSat and brain gray matter volume data;

[b] Numbers of subjects with available UrbanSat and behavioral data;

**Table S2. The correlation of UrbanSat with brain between-network FNC**

| Between network | CHIMGEN | | IMAGEN FU2 | | IMAGEN BL | |
|---|---|---|---|---|---|---|
| | *r* value | *P* value* | *r* value | *P* value* | *r* value | *P* value* |
| aDMN-ECN FNC | 0.165 | $3.25 \times 10^{-5}$ | 0.123 | $2.52 \times 10^{-3}$ | 0.258 | $1.16 \times 10^{-3}$ |
| aDMN-rFPN FNC | 0.133 | $8.34 \times 10^{-4}$ | 0.082 | 0.044 | 0.187 | 0.019 |
| aDMN-CBN FNC | 0.130 | $1.09 \times 10^{-3}$ | 0.086 | 0.035 | 0.162 | 0.043 |
| rFPN-LN FNC | 0.130 | $1.09 \times 10^{-3}$ | 0.087 | 0.033 | 0.194 | 0.015 |
| rFPN-lFPN FNC | 0.094 | 0.018 | 0.085 | 0.038 | 0.191 | 0.017 |
| AN-vSMN FNC | 0.131 | $1.04 \times 10^{-3}$ | 0.084 | 0.040 | - | - |
| AN-mVN FNC | 0.097 | 0.016 | 0.095 | 0.018 | - | - |

AN, auditory network; aDMN, anterior default mode network; BL, IMAGEN baseline assessment acquired at 14 years; CBN, cerebellar network; ECN, executive control network; FNC, functional network connectivity; lFPN, left frontal-parietal network; LN, language network; mVN, primary visual network; rFPN, right frontal-parietal network; vSMN, ventral sensor-motor network; FU2, IMAGEN follow up 2 assessment acquired at 19 years.

* *P* value after permutation test

**Table S3. Remote sensing satellite features information**

| Physical Properties | Data Sets | GEE ID | Time Span | Resource | Temporal Resolution | Spatial Resolution |
|---|---|---|---|---|---|---|
| GHSL | GHSL Population Grid (P2016) | JRC/GHSL/P2016/POP_GPW_GLOBE_V1 | 1975/1990/2000/2015 | Derived | epochs 4 time points | 250m |
| Night Lights | DMSP-OLS Nighttime Lights Time Series Version 4 | NOAA/DMSP-OLS/NIGHTTIME_LIGHTS | 1992.1.1-2014.1.1 | Derived | yearly | 30 arc-second (1km) |
| Climate | NOAA CDR AVHRR Normalized Difference Vegetation Index Version 4 | NOAA/CDR/AVHRR/NDVI/V4 | 1981.6.24-2017.10.05 | Derived | daily | 0.05°(5km) |
| Landsat | USGS Landsat 7 Collection 1 Tier 1 Raw Scenes | LANDSAT/LE07/C01/T1 | 1999.1.1-2017.10.08 | Raw | 16 days | 60 meters |
| Land Cover | Climate Change Initiative Land Cover datasets | - | 1992-2015 | Derived | - | 300 meters |

**Table S4. Landsat 7 band information**

| Name | Wavelength | Description (30m / pixel) |
|---|---|---|
| B1 | 0.45-0.52 um | Band 1 (blue) surface reflectance |
| B2 | 0.52-0.60 um | Band 2 (green) surface reflectance |
| B3 | 0.63-0.69 um | Band 3 (red) surface reflectance |
| B4 | 0.77-0.90 um | Band 4 (near infrared) surface reflectance |
| B5 | 1.55-1.75 um | Band 5 (shortwave infrared 1) surface reflectance |
| B6 | 10.40-12.50 um | Band 6 brightness temperature |
| B7 | 2.08-2.35 um | Band 7 (shortwave infrared 2) surface reflectance |

**Table S5. Remote sensing satellite features in the CHIMGEN and IMAGEN sample**

| Project | Satellite features | N | Time span | Mean (SD) |
|---|---|---|---|---|
| CHIMGEN | GHSL-POP | 3299 | 1990/2000/2015 | 14377.63（1265.72） |
| | Nighttime lights | 3301 | 1992-2014 | 28.69（18.35） |
| | NDVI | 3301 | 1987-2017 | 1333.63（304.18） |
| | Percentage of cropland | 3301 | 1992-2015 | 0.64（0.28） |
| | Percentage of urban build-up | 3301 | 1992-2015 | 0.29（0.26） |
| | UrbanSat | 3301 | - | 0.006（0.80） |
| IMAGEN | GHSL-POP | 1025 | 2000/2015 | 239.48（157.34） |
| | Nighttime lights | 1025 | 2009-2013 | 57.64（9.65） |
| | NDVI | 1025 | 2009-2015 | 1231.47（419.55） |
| | Percentage of cropland | 1025 | 2009-2015 | 0.20（0.23） |
| | Percentage of urban build-up | 1025 | 2009-2015 | 0.59（0.27） |
| | UrbanSat | 1025 | - | 0.04（0.86） |

GHSL-POP, Population density grid from Golbal Human Settlement Layer; NDVI, Normalized Difference Vegetation Index

**Table S6. Confirmatory factor analysis results**

| Sample | Factors | Year | CFI | AIC | BIC | [a]$R^2$ |
|---|---|---|---|---|---|---|
| CHIMGEN | 4 | 1990 | 0.983 | 4797.55 | 4832.00 | 0.758:0.693:0.908:0.366 |
| | 4 | 2000 | 0.991 | 30240.15 | 30288.92 | 0.790:0.655:0.890:0.244 |
| | 4 | 2015 | 0.999 | 31045.54 | 31094.36 | 0.720:0.672:0.765:0.296 |
| IMAGEN | 4 | 2000-2015 | 0.960 | 10148.502 | 10188.09 | 0.677:0.612:0.835:0.123 |

CFI, Comparative Fit Index; AIC, Akaike; BIC, sample-size adjusted Bayesian

[a] Ordered by percentage of urban build-up, percentage of cropland, nighttime lights and NDVI
